# A ligation-independent sequencing method reveals tRNA-derived RNAs with blocked 3’ termini

**DOI:** 10.1101/2023.06.06.543899

**Authors:** Alessandro Scacchetti, Emily J. Shields, Natalie A. Trigg, Jeremy E. Wilusz, Colin C. Conine, Roberto Bonasio

## Abstract

Despite the numerous sequencing methods available, the vast diversity in size and chemical modifications of RNA molecules makes the capture of the full spectrum of cellular RNAs a difficult task. By combining quasirandom hexamer priming with a custom template switching strategy, we developed a method to construct sequencing libraries from RNA molecules of any length and with any type of 3’ terminal modification, allowing the sequencing and analysis of virtually all RNA species. Ligation-independent detection of all types of RNA (LIDAR) is a simple, effective tool to comprehensively characterize changes in small non-coding RNAs and mRNAs simultaneously, with performance comparable to separate dedicated methods. With LIDAR, we comprehensively characterized the coding and non- coding transcriptome of mouse embryonic stem cells, neural progenitor cells, and sperm. LIDAR detected a much larger variety of tRNA-derived RNAs (tDRs) compared to traditional ligation-dependent sequencing methods, and uncovered the presence of tDRs with blocked 3’ ends that had previously escaped detection. Our findings highlight the potential of LIDAR to systematically detect all RNAs in a sample and uncover new RNA species with potential regulatory functions.

## Introduction

The assortment of RNA molecules present in prokaryotic and eukaryotic cells is vast, with sizes ranging from tens to thousands of nucleotides, and many different functionalities. One of the main criteria for RNA classification is based on their coding capacity, which distinguishes protein-coding RNAs (messenger RNAs; mRNAs) from non-coding RNAs (ncRNAs). The latter class can be further subdivided into structural ncRNAs, such as ribosomal RNAs (rRNAs), and transfer RNAs (tRNAs) that are required for translation; small nuclear RNAs (snRNAs) and small nucleolar RNAs (snoRNAs) that participate in splicing and rRNA biogenesis; and regulatory ncRNAs such as PIWI-interacting RNAs (piRNAs), micro RNAs (miRNAs), and long non-coding RNAs (lncRNAs) (Cech and Steitz, 2014; Eddy, 2001; Storz, 2002). In general, most regulatory ncRNAs are shorter than 50 nts and are often referred to as “small RNAs” (Ghildiyal and Zamore, 2009). Regulatory RNAs larger than 500 nts that do not code for proteins are referred to as long ncRNAs (lncRNAs) (Mattick et al., 2023).

A powerful approach to detect and quantify RNAs at the transcriptome-wide level consists of cloning them into cDNA libraries for next-generation sequencing (RNA-seq). Library construction techniques can be classified in three categories: mRNA-seq, Smart-seq, and small-RNA-seq. The most commonly used technique is mRNA-seq, whereby polyadenylated (polyA+) mRNAs are purified, fragmented, converted to cDNA via random-primed reverse transcription (RT), and then ligated to adapters for PCR amplification and sequencing (Mortazavi et al., 2008). These protocols have been standardized for abundant (> 500 ng) starting material and have also been expanded to non-polyA transcripts, typically by first removing rRNAs in place of the oligo(dT) based purification step (Hrdlickova et al., 2017; Yang et al., 2011). Smart-seq is one of several techniques based on template switching and Tn5-mediated tagmentation, which allow the construction of high-complexity libraries starting from limiting amounts of mRNA, including from single-cells (Hagemann-Jensen et al., 2020; Hagemann-Jensen et al., 2022; Hahaut et al., 2022; Picelli et al., 2013; Ramskold et al., 2012), which is not possible with traditional mRNA-seq. Neither mRNA-seq nor Smart-seq captures small RNAs, which are instead typically cloned into sequencing libraries by direct ligation of sequencing adapters to the 5’ and 3’ termini of the small RNA, followed by cDNA synthesis via RT (Baran-Gale et al., 2015; Dard-Dascot et al., 2018). While this methodology has been extremely successful, it can only capture RNAs with defined chemical structures at their termini, because the ligations can only proceed on RNAs that have a 5’ phosphate (5’P) and a 3’ hydroxyl (3’OH). Several ncRNAs, however, present chemical modifications at their termini (Crocker et al., 2022; Shi et al., 2022), posing a substantial challenge to standard ligation-based methods. Additional enzymatic steps and various strategies have been implemented to overcome this limitation (Behrens et al., 2021; Cozen et al., 2015; Gustafsson et al., 2022; Isakova et al., 2021; Mohr et al., 2013; Shi et al., 2021; Upton et al., 2021; Wang et al., 2021; Wulf et al., 2022; Xu et al., 2019; Zheng et al., 2015), but they require prior knowledge on the nature of chemical termini to be “repaired”, and, in some cases, suffer from substantial bias. Thus, an unbiased, ligation-independent method for small RNAs, ideally one that also captures longer RNAs, remains a critical need for the field.

Here, we describe a method to achieve ligation-independent detection of all types of RNA (LIDAR), regardless of their size or the chemical structure of their 5’ and 3’ termini. We combined a carefully designed template-switching oligo that contains unique molecular identifiers (UMIs) with quasi-random hexamer priming to minimize the formation of adapter dimers. This allowed us to avoid the final size-selection step of library construction, maximizing recovery of both small and long RNAs in the resulting libraries. Because LIDAR is ligation-independent, it captures RNA with modifications at their 3’ ends and retains the sensitivity of Smart-seq for low amounts of input RNA. LIDAR can be used as an efficient “all-in-one” method to analyze gene expression changes, with accuracy comparable to that of dedicated small and long RNA sequencing protocol. LIDAR captured RNA species from mouse embryonic stem cells and sperm that are notoriously difficult to clone by standard methods—including full-length tRNAs and derived fragments and, importantly, revealed the presence of 3’-blocked tRNA-derived RNAs (tDRs) that had escaped detection by sequencing until now.

## Results

### Development of LIDAR

We developed LIDAR with the goal of extending the ligation-independent nature of Smart-seq towards more RNA classes while retaining its high sensitivity for a broad range of transcripts in low-input conditions (Ramskold et al., 2012). All versions of Smart-seq, including the latest Smart-seq3 protocol (Hagemann-Jensen et al., 2020), rely on a template-switch strategy, which introduces sequences needed for library amplification at the 5’ end, bypassing the need to ligate a 5’ adapter to the RNA or cDNA. However, conventional Smart-seq was developed to sequence polyA+ RNAs and, therefore, it requires a defined sequence (polyadenylation) at the 3’ terminus. Modified versions of Smart-seq have been developed to expand the repertoire of clonable RNAs, but they rely either on *in vitro* polyadenylation (Isakova et al., 2021), which can only target RNAs with free 3’OH, or on random hexamer priming followed by size-selection of relatively large cDNAs to remove abundant adapter dimers, thus compromising the detection of RNAs smaller than 50 nts (Wang et al., 2023).

We introduced four key modifications to the Smart-seq3 protocol to extend the suitable substrates to all types of RNA, regardless of their size or 3’ terminal chemical structure (**Fig. 1A**). First, as a primer for RT, we utilized a “*quasi*-random” hexamer oligo devoid of cytosine at the terminal 3’ nucleotide (5’-NNNNND-3’; D = A, T, or G). This reduces the annealing of the RT primer oligo to the 3’-terminal GGG sequence of the template-switch oligo (TSO), therefore minimizing the generation of adapter dimers (**Fig. 1B**, **Fig. S1A**) (Ellefson et al., 2016; Seow et al., 2017). Second, we designed a TSO, with a sparse UMI structure, 5’-NNcgNNagNN-3’, preceding the 3’ terminal G’s, instead of the 5’-NNNNNNNN-3’ sequence of the original Smart-seq3 design (Hagemann-Jensen et al., 2020). This TSO design further reduces the formation of adapter dimers compared to the Smart-seq3 approach, favoring the generation of libraries with inserts originating from the input RNA (**Fig. 1B**). Third, we excluded the TSO from the first step of the reaction, thereby promoting the annealing of the RT primer to the RNA substrate and the binding of the RT enzyme to RNA–DNA hybrids (Arezi and Hogrefe, 2009). The RT reaction was allowed to proceed for 10 minutes at 25°C (**Fig 1A**, step I) and then raised to 42°C when the TSO was added to complete cDNA synthesis and template switching (**Fig. 1A**, step II). Fourth, we omitted the tagmentation and size selection steps after pre-amplification (**Fig 1A**, step III), proceeding directly to the final barcoding PCR (**Fig. 1A**, step IV). Together these modifications to the Smart-seq3 protocol allowed us to capture and sequence RNA of all sizes, including small RNAs in the 20–50 nts range.

**Figure 1.**
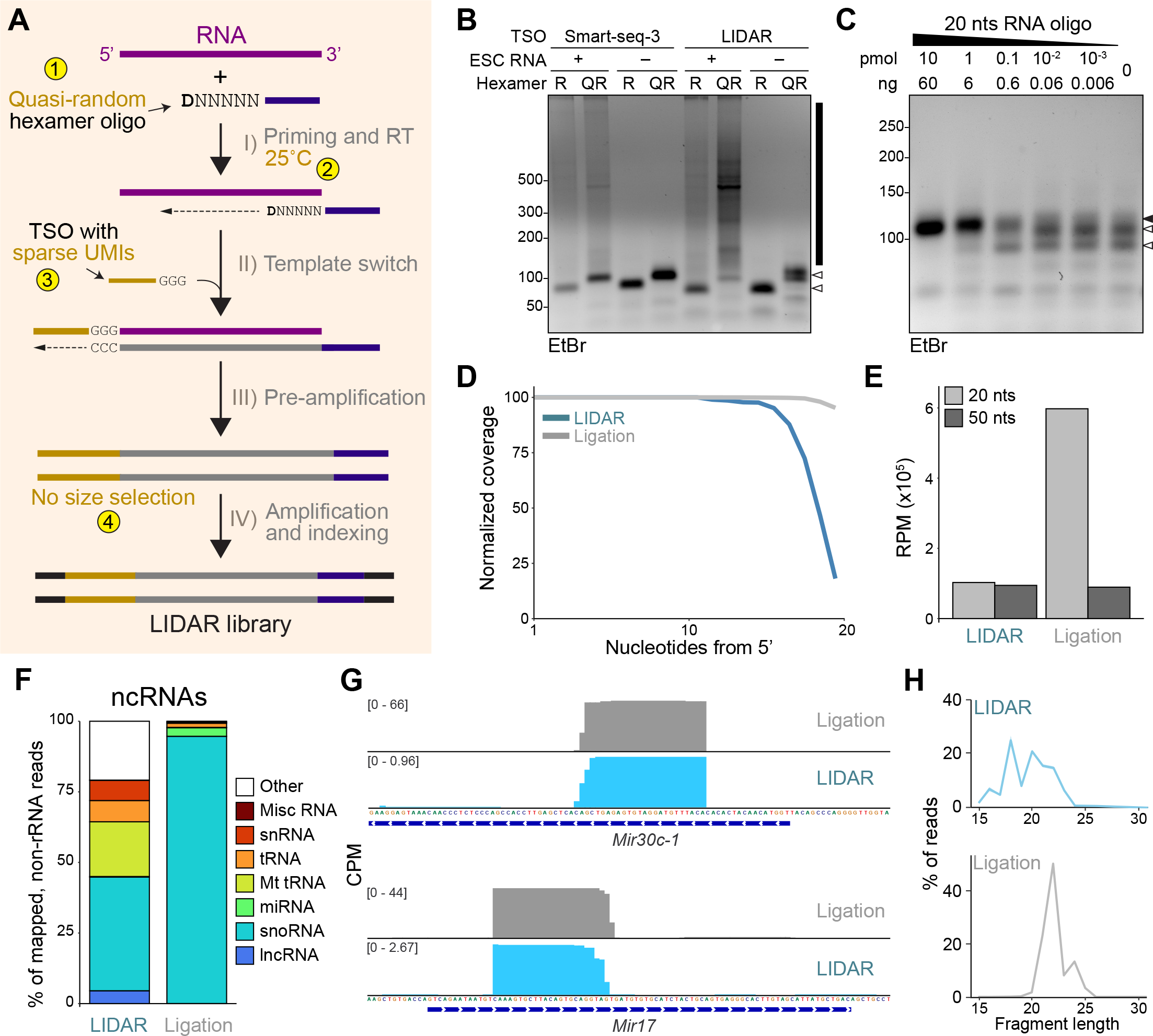
A modified Smart-seq3 protocol captures small RNAs (A) Schematic of LIDAR protocol. The four key modifications to the Smart-seq3 protocol are indicated with numbers highlighted in yellow. (B) Agarose gel electrophoresis of pre-amplification (step III) libraries constructed from total ESC RNA using the Smart-seq3 or LIDAR TSO, and random (R) or quasi-random (QR) RT primers. The black line on the side indicates productive libraries; the white arrowheads indicate adapter dimers. (C) Agarose gel electrophoresis of pre-amplified (step III), LIDAR libraries constructed from the indicated amounts of a synthetic 20 nts RNA. The black arrowhead indicates libraries with the 20 bp insert; the white arrowheads indicate adapter dimers. (D) Read coverage of a 20 nts RNA molecule cloned with LIDAR (blue) or with a conventional ligation-dependent protocol (gray). Read density is expressed as % of maximum coverage in each method. (E) Number of reads per million (RPM) sequenced mapping to 20 nts or 50 nts synthetic RNAs cloned by LIDAR or ligation. (F) Average (n = 3) biotype distribution of ncRNAs, expressed as % of mapped reads, in LIDAR and ligation-based libraries from total ESC RNA. (G) Genome browser snapshot of average LIDAR and ligation-based read coverage (expressed as counts per million, CPM) on two example miRNAs. (H) Average size distribution of reads mapping to miRNAs in LIDAR libraries starting from total ESC RNA (top) or ligation-based libraries (bottom). Data from 3 biological replicates.

### LIDAR captures small RNAs

Although priming with random hexamers is an extensively utilized method for cDNA synthesis (Hrdlickova et al., 2017), it has not been employed to clone and sequence small RNAs, likely due to concerns regarding efficiency of RT initiation on short templates, and to the difficulties in separating the resulting libraries with small inserts from empty libraries formed only by adapters. We reasoned that our use of template-switching technology combined with the suppression of adapter dimers would allow us to obtain LIDAR products from small RNAs. We tested LIDAR on synthetic RNAs of 20 nts and 50 nts that contained 8 random nucleotides at their 3’ end. Pre-amplification products of the correct insert size were obtained from as little as 0.6 ng of input RNA, indicating good sensitivity of LIDAR (**Fig. 1C**, **S1B**). Sequence coverage of both 20 nts (**Fig. 1D**) and 50 nts (**Fig. S1C**) RNAs by LIDAR was nearly complete, with 90% of the length of the oligos covered in > 70% of the aligned reads, indicating that, with appropriate modifications, an RT-based sequencing strategy can be used to capture small RNAs, similar to ligation-based approaches. In fact, even a ligation-based approach (NEB kit, see Experimental Procedures; henceforth also referred to as “ligation”) did not result in 100% coverage at the 3’ (**Fig. 1D, S1C**), likely due to incomplete synthesis or partial degradation of the RNA oligonucleotides.

Older ligation-based cloning methods displayed substantial preference for certain sequences in the target RNA, especially at its 3’ end (Raabe et al., 2014). LIDAR showed no bias in the sequence of the substrate RNA (**Fig. S1D**, top). The ligation-based protocol we employed also displayed a very minimal bias in our hands (**Fig. S1D**, bottom), indicating that modern versions of this experimental approach have successfully addressed this issue. When an equimolar mix of 20 nts and 50 nts oligos was used as input, LIDAR captured them equally well, whereas the ligation-dependent method preferentially recovered the smaller RNA, consistent with the fact that this protocol was optimized for sequencing canonical small RNAs such as miRNAs (**Fig. 1E**).

Because LIDAR captured artificial RNAs as short as 20 nts *in vitro*, we wondered if the approach could capture endogenous small RNAs within total RNA extracted from mouse embryonic stem cells (ESC). We successfully constructed LIDAR libraries from small amounts of total cellular RNA, as little as 1 ng (**Fig. S1E**), although with increasing contamination of empty libraries from adapter dimers (**Fig. S1F**). cDNA synthesis events from the TSO directly annealing to RNA (TSO strand invasion events), measured by counting reads mapping to genomic loci preceded by their matched UMI sequence, were also limited (**Fig. S1G**), occurring at a frequency comparable to FLASH-seq, a method developed to reduce strand invasion events in Smart-seq3 (Hahaut et al., 2022).

Because LIDAR is designed to capture all types of RNA, we expected a large proportion of reads to map to abundant structural RNAs, such as ribosomal RNA (rRNA). In fact, 64% of LIDAR reads mapped to rRNA (**Fig. S1H**), three times more than with the traditional ligation protocol (**Fig. S1H**) but similar to ligation-based techniques aimed at capturing comprehensive sets of RNAs, such as PANDORA-seq (Shi et al., 2021). Analysis of ncRNAs detected by LIDAR revealed broad representation of several short RNA biotypes, such as snoRNA, snRNAs, and mitochondrial/cytosolic tRNAs, while libraries constructed with the ligation protocol were largely comprised of snoRNAs (**Fig. 1F**). Reads from LIDAR libraries also mapped to mature miRNAs (**Fig. 1G**), and despite their small size (20–24 nts), their entire length was often covered (**Fig. 1G–H**). As expected, capture of miRNAs by LIDAR was less efficient compared to ligation-based methods specifically optimized for these small RNAs. However, we were able to improve their representation in LIDAR libraries by size-selecting the input for RNAs < 200 nts or < 50 nts by silica column or PAGE purification, respectively (**Fig. S1I-K**).

Overall, our data show that LIDAR is a versatile, ligation-independent method that allows cloning and sequencing of small RNAs of any size and biotype from small amounts of input material, requiring less than 4 hours to complete, with minimal hands-on time.

### Simultaneous detection of differentially expressed protein-coding and small RNAs

Because the LIDAR protocol was designed to capture RNA of all sizes, including small RNAs, we omitted the tagmentation step of Smart-seq3, to avoid their fragmentation into unclonable and unmappable fragments. Without tagmentation, full-length cDNAs from long (> 500 nts) transcripts would not be sequenced efficiently on the Illumina platform (Tan et al., 2019): however, the *quasi-*random hexamers should prime RT from multiple sites within a long transcript and, therefore, all transcripts regardless of size should be detected in the final libraries. The large representation of reads from rRNA (**Fig. S1H**) suggested that this was the case.

After excluding those mapping to rRNA, a large proportion of the remaining reads originated from protein-coding mRNAs, which were virtually undetected by the ligation-based method (**Fig. S2A**). Consistent with this result, LIDAR libraries contained inserts from a much larger range of original transcript size (**Fig. S2B).** We sought to determine if this extensive coverage would allow LIDAR to detect differentially expressed genes of various biotypes and sizes. We utilized an established *in vitro* differentiation system in which ESC are converted to neural progenitor cells (NPC)—a process accompanied by dramatic changes in gene expression profiles (Gouti et al., 2014; Petracovici and Bonasio, 2021) (**Fig. 2A**). Using conventional mRNA-seq (polyA+ purification followed by chemical fragmentation and random-primed RT), we detected 8,955 genes differentially expressed (adjusted *p* < 0.05) between NPC and ESC (**Fig. 2B,** left). These included pluripotency markers (*Sox2, Nanog*, *Pou5f1*) downregulated upon differentiation, and neuronal markers (*Nestin, Ncam1, Neurog1*) upregulated in NPC. LIDAR libraries from total RNA revealed a comparable number of differentially expressed protein-coding genes (*n* = 5,182; adjusted *p* < 0.05) (**Fig. 2B**, right), including the same known markers of pluripotency and differentiation. Comparison of the differentially expressed genes detected by mRNA-seq and LIDAR showed very similar expression patterns (**Fig. 2C**), a high degree of overlap (**Fig. 2D**), and a good correlation between fold-changes (**Fig. 2E**). Despite the fact that LIDAR detected overall fewer miRNAs, they were quantified accurately, as the profiles of differentially expressed miRNAs between ESC and NPC were in good agreement with those obtained with a ligation-based cloning method (**Fig. 2F–H**). Correlated results, although to a lesser extent, were observed for the snoRNA and snRNA profiles (**Fig S2C–F**).

**Figure 2.**
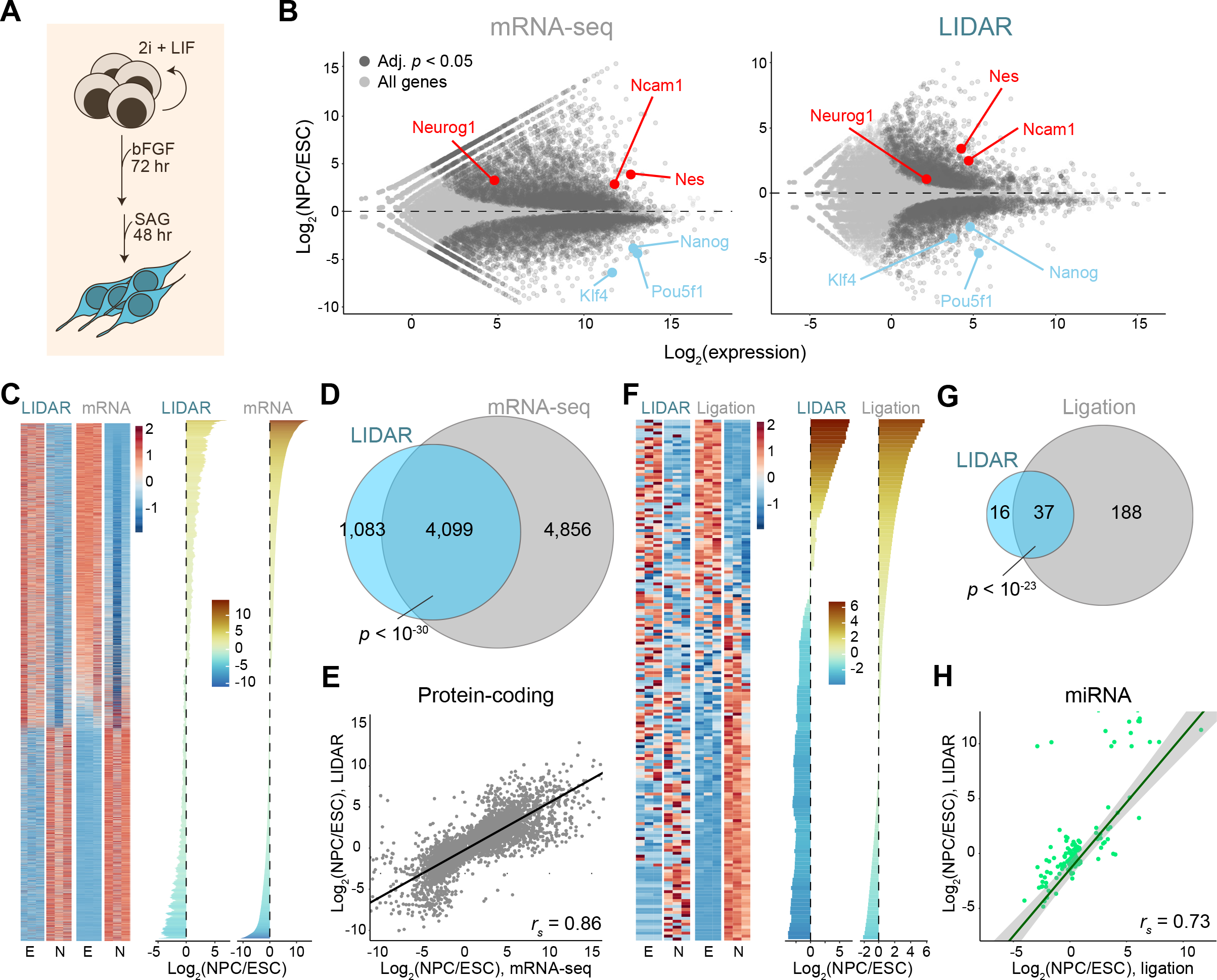
**LIDAR detects differential expression in small and large RNAs** (A) Schematic of ESC to NPC differentiation protocol. (B) MA plot of gene expression changes between NPC and ESC measured by mRNA-seq (left panel) or LIDAR (right panel) from total RNA. Dark grey, genes with significant changes (adjusted p value < 0.05, n = 3). Red, NPC markers. Blue, ESC markers. (C) Heatmap of z score-converted expression levels, calculated as transcripts per million (TPM), of differentially expressed protein-coding genes between NPC (N) and ESC (E), as measured by LIDAR or mRNA-seq. Individual replicates are shown. Rolling mean of log2 fold-change (n = 3) is shown on the right. (D) Venn diagram for differentially expressed protein-coding genes detected in LIDAR (blue) or mRNA-seq (grey). The p value for the overlap is from a hypergeometric test. (E) Correlation of average log2 fold changes (NPC vs. ESC, n = 3) of differentially expressed protein-coding genes detected by LIDAR and mRNA-seq. rs, Spearman’s rank correlation coefficient. (F) Heatmap as in (C) for miRNAs comparing LIDAR or ligation-based libraries starting from total RNA. (G) Venn diagram for differentially expressed miRNAs detected in LIDAR (blue) or ligation-based libraries (grey). The p value for the overlap is from a hypergeometric test. (H) Correlation plot as in (E) for differentially expressed miRNAs detected by LIDAR vs. ligation-based libraries.

These results highlight the effectiveness of LIDAR in analyzing gene expression changes simultaneously in distinct RNA populations, which traditionally would have been measured in separate experiments with different library construction protocols.

### LIDAR reveals 30–40 nt 3’ tDRs in ESC

tRNA-derived RNAs (tDRs) (Holmes et al., 2023) constitute an emerging class of small ncRNAs with a growing list of functional roles in the regulation of translation (Kim et al., 2017), transposons (Schorn et al., 2017), and, possibly, transgenerational epigenetics (Chen et al., 2016; Sharma et al., 2016). Multiple classes of tDRs have been recognized, based on their size and position within the tRNA sequence (**Fig. 3A**) (Su et al., 2020; Xie et al., 2020). 5’ tDRs start at the 5’ position and end either at or right after the D loop (5’-tRF), or at the anticodon (AC) loop (5’ tRNA halves, also known as 5’-tiRNA). 3’ tDRs end at the 3’ end of mature tRNAs, including the non-templated CCA sequence, and start at the T-loop (3’-tRF) or at the AC-loop (3’ tRNA halves, also known as 3’-tiRNA). Other tDRs that do not start at the 5’ end or do not end at the 3’ end are classified as internal tDRs (i-tRF).

**Figure 3.**
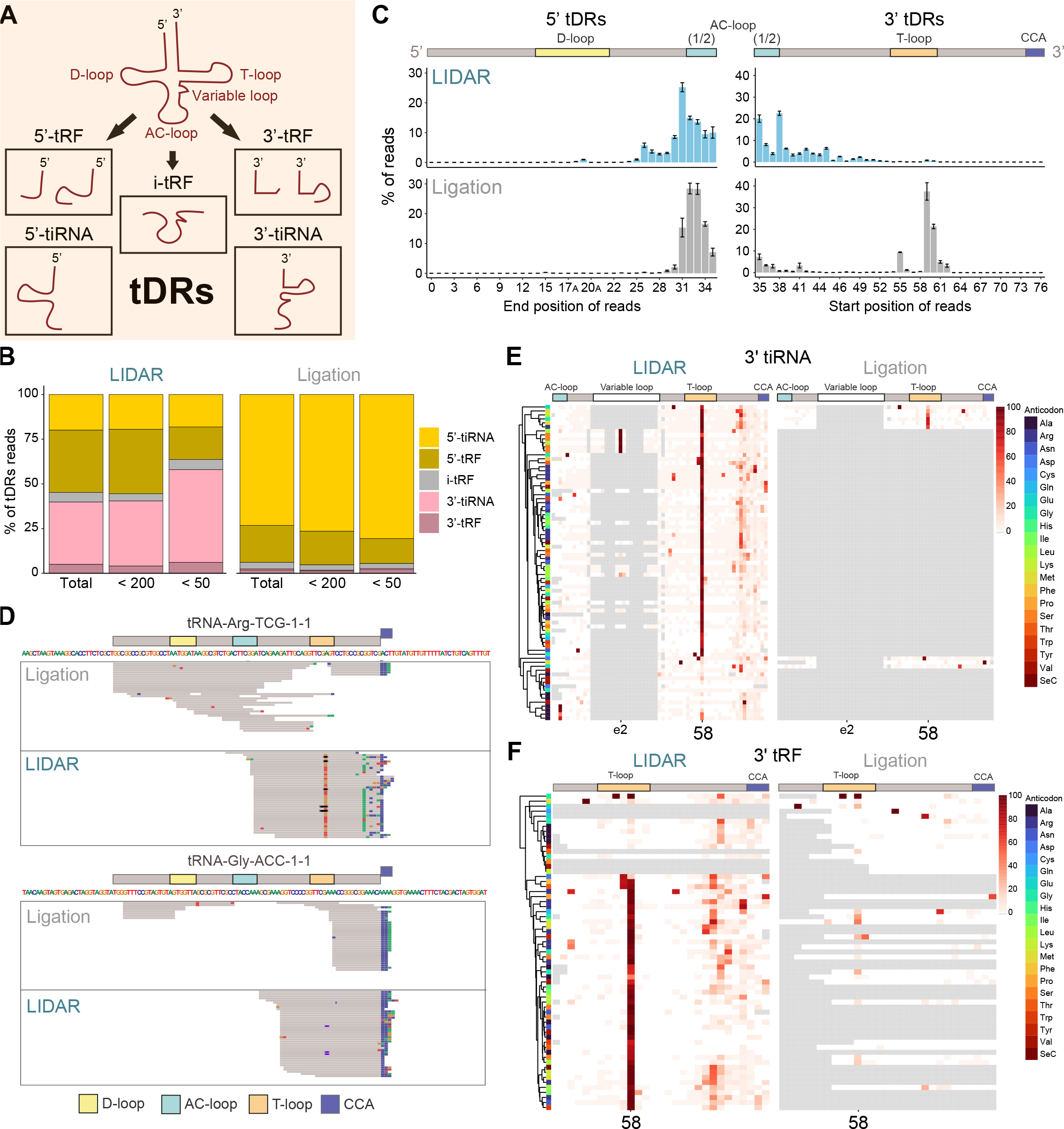
**LIDAR uncovers 3’ tDRs not efficiently captured by ligation** (A) Scheme of possible tRNA-derived RNAs (tDRs). (B) Average (n = 3) tDRs distribution, expressed as % of reads mapping to any tDR, in LIDAR and ligation-based libraries from total, < 200 nts, and < 50 nts ESC RNA. (C) Histogram of average (n = 3, ⊠ SEM) read end (left) or beginning (right) position frequency, expressed as % of all reads mapping to the corresponding tRNA fragment type, for 5’ tDRs (tRF or tiRNA) (left) and 3’ tDRs (tRF or tiRNA) (right) in LIDAR (blue) or ligation-based (grey) libraries starting from < 200 nt ESC RNA. (D) Example genome browser snapshots showing single collapsed mapped reads mapping to two different tRNAs in LIDAR or ligation-based libraries starting from < 200 nt ESC RNA. Position of various loops, as shown in Fig. 3A, are indicated on top of each panel. The non-templated 3’-terminal CCA sequence is depicted as a blue box. (E) Heatmap of average misincorporation rate (expressed as % detected) for every canonical position (column) in every 3’-tiRNA iso-acceptor (rows) in LIDAR (left) or ligation-based (right) libraries starting from < 200nt ESC RNA (n = 3). In gray, positions with coverage = 0. Position of loops, as shown in Fig. 3A, are indicated on top. (F) Same as (E) but for 3’-tRF.

Ligation-based libraries are strongly biased towards the detection of 5’ tRFs and 5’-tiRNAs and do not capture efficiently 3’ tRFs and 3’-tiRNAs, despite the fact that they can be detected in northern blots (Gustafsson et al., 2022; Sharma et al., 2018). The reasons for this bias are unclear. We speculated that chemical modifications at the 3’ end might interfere with ligation and wondered whether LIDAR would capture tDRs that cannot be cloned by ligation-dependent methods. In libraries constructed from size-selected ESC RNA smaller than 200 nts, we detected a far greater proportion of reads mapping to 3’ tDRs in LIDAR compared to ligation-dependent libraries, which, as previously reported, showed a strong bias for 5’ fragments (**Fig. 3B**) (Gustafsson et al., 2022). The size distribution for reads mapping to the 5’ portion of tRNA gene models was similar in both library construction methods (**Fig. 3C**, left), suggesting that the majority of LIDAR reads assigned to this class originated from actual 5’ tDRs. On the other hand, the size distribution of 3’ fragments obtained by LIDAR was markedly different compared to that observed in ligation-based libraries (**Fig. 3C–D**). In LIDAR, the majority of reads mapping to the 3’ of tRNA genes were between 30 and 40 nts in length, whereas the 3’-tRFs captured by ligation were mostly 17–22 nts, corresponding to cleavage events within the D-loop (**Fig. 3C**, right) (**Fig. S3A**). 3’ tDRs of size between 30 and 40 nts in length have been observed by northern blots across different human tissues and cell lines (Kawaji et al., 2008), including a ∼40 nt 3’ tRNA fragment from *ArgTCG-1-1* (Torres et al., 2019), with the same size as the one we identified with LIDAR (**Fig. 3D**). This suggest that our method can identify *bona fide* tDRs that escape detection in ligation-based methods.

While it is still unclear how different tDRs are generated, it is generally believed that many derive from enzymatic cleavage of mature tRNA molecules (Su et al., 2020). During their biogenesis, tRNAs undergo extensive chemical modification at stereotypical nucleotide positions, which are necessary for their function (Suzuki, 2021). Sites of tRNA modifications can be an obstacle for reverse transcriptases, resulting in mismatches or deletions during cDNA synthesis that can be utilized as indirect readouts to map modified residues (Ryvkin et al., 2013). Both 3’ tiRNA and tRFs showed high frequency of mismatches at position 58 in LIDAR (**Fig. 3D–F**), consistent with the presence of m1A (Behrens et al., 2021; Gogakos et al., 2017; Suzuki, 2021), which is typically found on mature tRNA and thought to have a stabilizing effect on their structure (Zhang and Jia, 2018). Thus, the 3’ tDRs detected by LIDAR may derive from mature and modified tRNAs by cleavage. Alternatively, the 3’ tDRs could be directly targeted by the RNA modification machinery. In ligation-based libraries, fewer 3’ tDRs with mismatches at position 58 were detected (**Fig. 3D–F**). Since the reverse transcriptase used in the ligation protocol is of the same family as the one used in LIDAR (M-MuLV), it is unlikely that the presence of m1A affects cDNA synthesis from 3’ tDRs in the ligation protocol, thus suggesting another determinant is responsible for the observed difference in 3’ tDRs representation between LIDAR and ligation-based libraries.

In many cases, the LIDAR reads mapping to the 3’ portion of tRNAs contained the non-templated CCA (**Fig. 3D**), further indicating that they originate from cleavage or processing products of mature tRNAs (Rubio Gomez and Ibba, 2020). However, in some cases the terminal CCA sequence was missing or incomplete (**Fig. 3D**), likely due to internal priming. This prompted us to develop a modified LIDAR protocol to increase 3’ end coverage. We designed an alternative RT strategy, whereby a fully random hexamer protrudes as a 3’ overhang of a double-stranded DNA molecule (**Fig. S3B**). The presence of the double-stranded stretch should constitute a steric hindrance to internal priming, favoring the initiation of RT at the 3’ end of the RNA substrate. When we constructed libraries with this modified LIDAR protocol (3’-LIDAR) using 20 nts and 50 nts synthetic RNAs as input, we observed an increased number of reads covering their entire sequences, corresponding to more priming events close to their 3’ (**Fig. S3C**). Consistent with this, 3’-LIDAR improved the coverage of the 3’ ends of tRNAs, resulting in more efficient capture of fragments ending in 3’ non-templated CCA (**Fig. S3D–E**), although at the expense of increased rates of adapter dimer formation (**Fig. S3F**). Size distribution and representation of 3’ tDRs were similar in LIDAR and 3’-LIDAR (**Fig. S3G–H**), further indicating that the unexpected 30–40 nts 3’ tDRs identified in LIDAR originated from mature, CCA-containing tRNAs.

In conclusion, LIDAR revealed 3’ tDRs that are longer than those cloned by ligation methods and that contained m1A, likely derived from cleavage of mature tRNAs.

### LIDAR clones charged full-length tRNAs and 3’ tDRs

One of the motivations to develop LIDAR was to allow the cloning and sequencing of RNAs with 3’ ends blocked by chemical modifications that make them inaccessible to conventional ligation-dependent protocols. The best known RNAs with these features are the U6 snRNA and tRNAs charged with amino acids (**Fig. 4A**). U6 is a major component of the spliceosome (Matera and Wang, 2014) and, unlike other snRNAs, it is transcribed in mammals by RNA Pol III and its 3’ terminus is processed to a 2’-3’-cyclic phosphate (2’-3’cP) (Didychuk et al., 2018). This 3’ end modification, unless resolved by T4 PNK treatment, prevents the ligation of a 3’ adapter (Shi et al., 2021). A similar impediment to ligation can arise from the aminoacyl moiety which, in a subset of tRNAs, is attached to the 3’ hydroxyl group by aminoacyl-tRNA synthetases (Rubio Gomez and Ibba, 2020). These can only be removed by chemical de-acylation or β-elimination (Evans et al., 2017; Shigematsu et al., 2017). Random-hexamer priming is based on hybridization, and therefore should not be affected by the chemical status of the 3’ of the RNA (**Fig. 4A**). Indeed, LIDAR successfully cloned a synthetic RNA with its 3’ hydroxyl group blocked by a biotin moiety, which was completely missed by ligation-based libraries (**Fig. 4B**, **S4A**).

**Figure 4.**
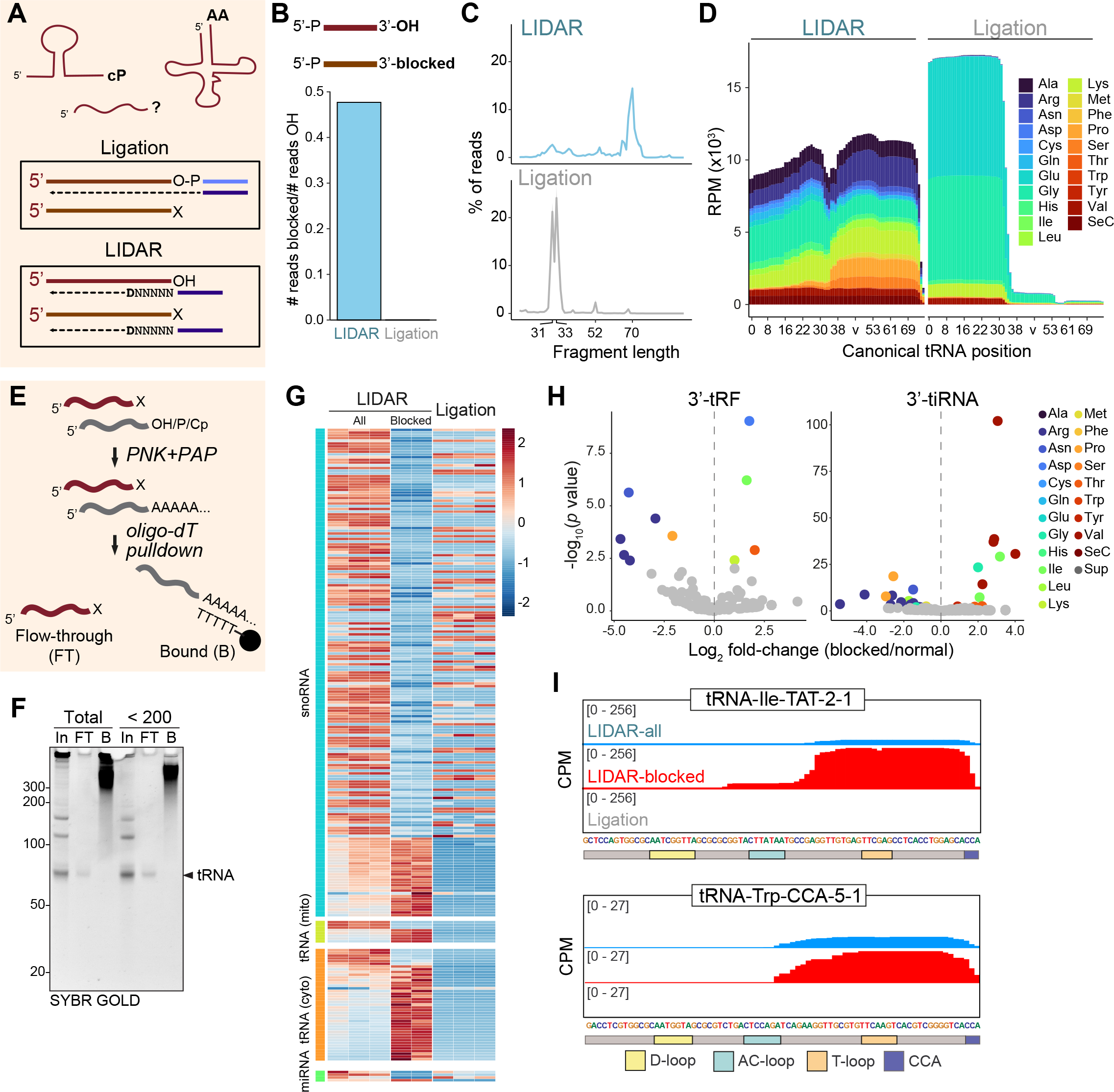
**LIDAR detects full-length tRNAs and tDRs with blocked 3’ ends** (A) Top: schematic of RNAs with possible blocked 3’ ends (Cp: 2’-3’ cyclic phosphate; AA: amino acid; ?: unknown). Bottom: only RNAs (red) with free 3’ ends can be efficiently cloned by both ligation-based protocols and LIDAR. RNA with blocked 3’ ends can only be captured by LIDAR. (B) Barplot of reads per million (RPM) ratio of sequences mapping to 3’ biotinylated (blocked) vs. 3’OH synthetic 20 nts RNA in LIDAR and ligation libraries. (C) Average (n = 3) size distribution of all reads mapping to tRNAs, expressed as a % of all tRNA reads, in LIDAR (blue, top) or ligation-based (grey, bottom) libraries from ESC RNA < 200 nts. (D) Average coverage and representation of ESC tRNA anticodons (color-coded) in LIDAR (left) or ligation (right) libraries from ESC RNA < 200 nts. v, collapsed variable loop position. Data from 3 biological replicates. (E) Scheme for the enrichment of RNA with blocked 3’ ends. Input RNAs with 3’OH, 3’P, and 2’-3’cP (cP) were end-repaired with T4 PNK and polyadenylated using E. coli PAP. The artificially polyadenylated RNAs were removed via oligo-dT beads (B). RNA with blocked 3’ end were not polyadenylated and remained unbound in the flow-through (FT). (F) Urea PAGE of total and < 200 nts ESC blocked RNA (FT) isolated using the method represented in Fig. 4E. 100% Input (In) and polyA bound (B) fractions loaded as controls. (G) Heatmap of z-score normalized TPM expression of differentially expressed (adj. p < 0.05) snoRNA, miRNA, and tRNA between LIDAR libraries from all and blocked ESC total RNA. Z-score normalized expression in ligation-based libraries from unfractionated (all) ESC RNA is also shown. (H) Volcano plot showing log2 fold-change in frequency, calculated over all reads mapping to tDRs of 3’-tRF and 3’-tiRNA isoencoders between LIDAR libraries from all < 200 nt RNA (n = 3) vs blocked (n = 2) RNA from ESC. Comparisons with adjusted p value < 0.05 are colored according to their anticodon. (I) Read density of LIDAR (from all or blocked ESC RNA < 200 nts) and ligation-based libraries on two example tRNAs.

Given that LIDAR captured a synthetic RNA with a blocked 3’ end, we reasoned that one cause for RNAs being captured by LIDAR but not ligation may be a 3’ block. Accordingly, LIDAR efficiently captured the U6 snRNA, which was virtually absent from ligation-based libraries, likely because its terminal 2’-3’cP impedes ligation (**Fig. S4B**). Furthermore, analysis of reads mapping to tRNA revealed a peak at ∼70 nts, which is almost absent in ligation-based libraries and similar to the size of mature tRNAs (**Fig. 4C**). This difference was not due to the bias of ligation-dependent libraries toward smaller inserts, since the larger snoRNAs were efficiently captured by ligation (**Fig. S4C**). LIDAR reads were distributed across tRNAs with a broad range of anticodons, similar to techniques developed specifically to clone mature tRNAs (Behrens et al., 2021) (**Fig. 4D**). In contrast, the ligation-dependent libraries were strongly biased towards a subset of anticodons (**Fig. 4D**), as recently reported (Gustafsson et al., 2022). Full-length tRNA reads from LIDAR libraries contained sequence mismatches at sites where mature tRNAs are known to be chemically modified, including position 9, 26, 32, 37, e2 (within the variable loop), and 58, similar to observations made with the dedicated mim-tRNAseq technique (Behrens et al., 2021) (**Fig. S4D,** left). Some of these modification sites were also detected as mismatches in reads from ligation-based libraries, but with much lower frequency (**Fig. S4D,** right). Given that typically more than 80% of mature cytosolic tRNAs are charged (Evans et al., 2017), we conclude that LIDAR captured those species much more efficiently than the ligation-based method, due to its ability to bypass blocked 3’ ends.

To better profile molecules with inaccessible 3’ ends, we depleted transcripts containing 3’ OH end with a biochemical method: we treated ESC RNA with PNK to convert 3’P and 2’-3’cP to 3’OH, added a polyA tail to all RNAs with a 3’OH, and then removed them by hybridization with oli-go-dT-conjugated magnetic beads (**Fig. 4E**). After magnetic separation, the flow-through is enriched for RNAs that do not have a 3’OH, 3’P or 2’-3’cP, i.e. they have a “blocked” 3’. The main RNA species in the blocked RNA preparation appeared as a band of ∼70 nts, corresponding to the size of mature tRNAs (**Fig. 4F**). Accordingly, reads from LIDAR libraries constructed on blocked RNAs mapped more frequently to tRNA genes (**Fig. S4E**), and were specifically enriched for full-length tRNA reads compared to untreated RNA inputs (**Fig. S4F**). Size distribution of reads mapping to tRNAs also showed a peak at ∼70nt for LIDAR, which increased when blocked RNA was used as input (**Fig. S4G**). LIDAR libraries from blocked RNAs were enriched for reads mapping to tRNAs, which were poorly represented in ligation libraries (**Fig. 4G**). Other small RNA classes, with the exception of a subset of snoRNAs, were captured efficiently in LIDAR libraries from all (i.e. non-blocked) RNA and also detected by ligation (**Fig. 4G**). Interestingly, we found several 3’ tDRs enriched in the blocked RNA population (**Fig. 4H–I**). We speculate that at least some of these 3’ tDRs may be aminoacylated, thus deriving from cleavage of mature and charged tRNAs. A recent report provided the first evidence of aminoacylated 3’ tDR (Liu et al., 2021), supporting the existence of a new 3’ tDR class that can now be detected with LIDAR.

Thus, LIDAR is effective in cloning RNAs with blocked 3’ and can be used to analyze RNA populations that are not detectable using ligation-based methods, including fulllength aminoacylated tRNAs and their 3’ fragments.

### LIDAR captures transcript diversity in mouse sperm

Ligation-based small RNA sequencing studies reported that the RNA content of mouse sperm purified from the cauda (distal) region of the epididymis is dominated by small- and medium-size species (< 2 kb), in particular 5’ tDRs (Chen et al., 2016; Sharma et al., 2016). However, analyses of sperm RNA content by northern blot or the recently described OTTR-seq (Gustafsson et al., 2022) revealed the presence of 3’ tDRs—and possibly also full-length tRNAs—that were missed by conventional small RNA library preparation protocols (Sharma et al., 2018). We thus tested whether LIDAR could be employed to generate a more comprehensive catalog of the RNA content of sperm.

RNAs isolated from cauda sperm were predominantly small (**Fig. S5A**), as previously reported (Gustafsson et al., 2022; Sharma et al., 2018; Shi et al., 2021). We prepared LIDAR and ligation-based libraries from either untreated sperm total RNA or after enriching for blocked transcripts, as above (**Fig. 4E**). As control, we also sequenced libraries obtained by ligation. As in the case of ESC, many LIDAR reads mapped to rRNA (71%) (**Fig. S5B)**, almost three times higher than in ligation libraries (28%), likely because rRNA fragments in sperm have 3’P or 2’3’cP ends that require enzymatic conversion before ligation, as shown with PANDORA-seq (Shi et al., 2021). LIDAR libraries from blocked sperm RNA also contained a large proportion of rRNA reads (71%). While it is possible that our biochemical depletion was incomplete, we noticed an enrichment of blocked reads mapping to well-defined regions of 18S and 28S rRNAs (**Fig. S5C**), suggesting the possibility that previously unreported small rRNA fragment might be chemically blocked at their 3’ ends. As in the case of ESC, LIDAR captured a higher amount of U6 (**Fig. S5D**), indicating that sperm may also contain mature U6 RNA molecules, possibly ending with a 2’-3’cP.

After excluding rRNAs, the majority of the RNA fragments cloned from cauda sperm by both LIDAR and ligation originated from tRNAs (**Fig. 5A**), in line with previous studies (Chen et al., 2016; Sharma et al., 2016; Sharma et al., 2018). Nonetheless, we observed several differences between LIDAR and ligation libraries. Both the cytosolic and the mitochondrial tRNA pools were well represented in LIDAR, whereas the ligation-based protocol favored cytosolic tRNAs almost exclusively (**Fig. 5A**). The size distribution of cytosolic tRNA reads revealed that a large fraction originated from tDRs rather than full-length mature tRNAs, as previously reported (Sharma et al., 2018) (**Fig. 5B**). On the other hand, reads mapping to mitochondrial tRNAs peaked at ∼70 nts in LIDAR libraries and increased in libraries enriched for blocked RNAs (**Fig. 5A–B**). This indicates that the mitochondrial tRNAs in sperm are 1) protected from fragmentation and 2) blocked at their 3’ end, likely by their amino acid. This also demonstrates that the shorter fragments mapping to cytosolic tRNAs detected in these samples constitute *bona fide* RNA species and are not LIDAR artifacts.

**Figure 5.**
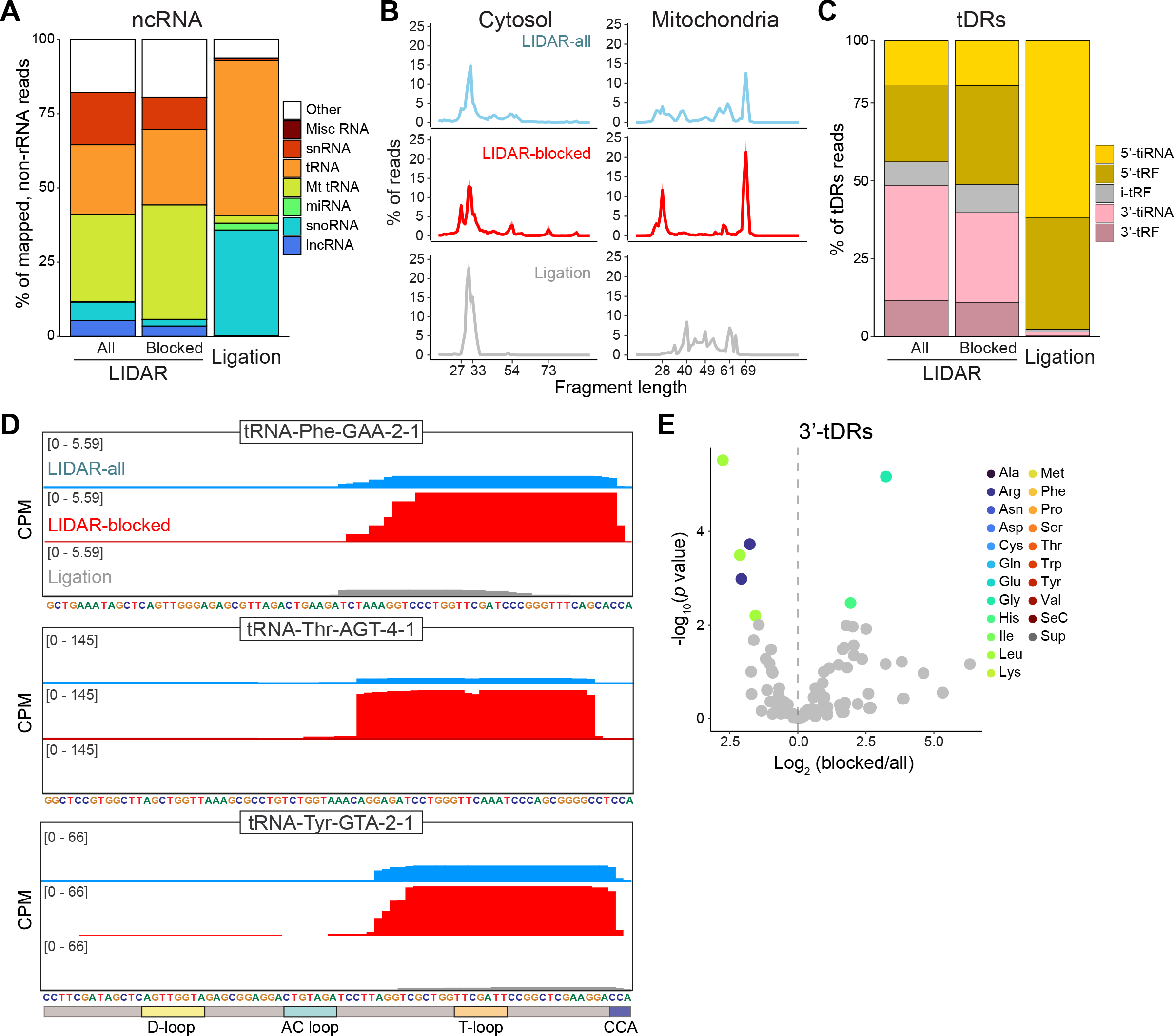
**Sperm contains full-length tRNAs and 30–40 nts 3’ tDRs** (A) Average (n = 2) biotype distribution of ncRNAs, expressed as % of mapped reads, in LIDAR from sperm RNA before or after enriching for blocked RNA and in ligation-based libraries. (B) Average (n = 2) size distribution of reads mapping to cytosolic or mitochondrial tRNAs, expressed as % of reads of all tRNA reads. Libraries obtained with LIDAR from all or blocked sperm RNA are shown as well as ligation-based libraries. (C) Average (n = 2) tDR distribution, expressed as % of reads mapping to any tDR, in LIDAR from all sperm RNA, blocked RNA, or ligation-based libraries. (D) Read density of LIDAR (from all or blocked sperm RNA) or ligation-based libraries on three example tRNAs. (E) Volcano plot showing log2 fold-change for LIDAR libraries on blocked vs. all RNA from sperm of individual 3’ tDRs (tRFs and tiRNAs) from isoencoders. Fragments for which the comparison has an adjusted p value < 0.05 are colored according to their anticodon.

As we observed in ESC, 5’ and 3’-derived tDRs were detected at similar levels in LIDAR libraries, whereas ligation favored 5’ tDRs (**Fig. 5C**). The few 3’ tDRs detected by ligation-based libraries were mostly small (17–22 nts), although larger species were also present (**Fig. S5E**). LIDAR, as in ESC, captured mostly fragments 30–40 nts in size (**Fig. S5E**). Importantly, LIDAR identified several 3’ tDRs that were not detected or were very underrepresented in ligation libraries (**Fig. 5D**), contained a heavily modified base at position 58 (**Fig. S5F**), and were enriched in the blocked RNA population (**Fig 5E**, **S5F**), suggesting that members of the new class of long 3’ tDRs with blocked 3’ ends that we detected in ESC are also present in sperm.

In summary, LIDAR uncovered an unexpected diversity in the sperm RNA payload, including full length and 3’ tDRs that had previously escaped detection.

## Discussion

Here, we presented LIDAR, a new RNA-seq technique that utilizes a custom template-switch strategy combined with *quasi-*random hexamer priming to sequence all types of RNA, regardless of size or chemical modifications at 5’ and 3’.

### Technical advantages of LIDAR

LIDAR enables quick and comprehensive coding and non-coding transcriptome analysis. The protocol (**Fig. 1A**) can be completed in less than 4 hours, faster than the most recent methods based on template switching (Hagemann-Jensen et al., 2022; Hahaut et al., 2022; Isakova et al., 2021), and with minimal hands-on time. It does not require a tagmentation step and all molecules contain UMIs, which can increase quantification accuracy, especially from low-input samples, and are missing from most commercial and non-commercial small RNA cloning strategies. Additionally, the sparse UMI structure allows the sequencing of LIDAR-seq libraries without major issues in cluster detection during Illumina sequencing caused by low complexity at the 5’ end of libraries (Krueger et al., 2011; Wu et al., 2015).

The input RNA for LIDAR does not require chemical or enzymatic pre-processing to remove 3’ terminal modifications. This confers two key advantages to LIDAR: 1) it minimizes sample loss due to RNA degradation and contamination during pre-processing steps, and 2) it bypasses the requirement of *a priori* knowledge of terminal RNA modifications.

LIDAR can detect transcriptional changes of most RNAs simultaneously (**Fig. 2**), with performance comparable to tools specifically dedicated to the analysis of certain RNA populations. In the case of the *in vitro* neural differentiation model we used for our analysis, the profile of changes in protein-coding RNAs and small RNAs (e.g.: miRNAs), defined by specialized protocols, were well recapitulated by LIDAR. LIDAR requires relatively low input amounts (10– 500 ng total RNA), making it amenable for comprehensive transcriptome analyses of biological sources where starting material is limited, such as sorted cells or tissue biopsies.

We also developed 3’-LIDAR (**Fig. S3**), whereby the forced annealing of the random hexamer to the 3’ end allowed us to obtain better coverage of the 3’ terminal portions of the RNA inserts.

### Ligation-independent small RNA sequencing

The dependence on ligation in the majority of small RNA cloning protocol requires free 3’OH ends on the target RNA or a means to generate them. As an example, the recently developed PANDORA-seq method utilizes a combination of enzymatic treatments to generate RNA ends compatible with ligation as well as removal of internal RNA modification that could stall RT enzymes (Shi et al., 2021). Adaptations of template switching protocols that include an oligo-dT priming step have been introduced to allow small RNA detection by means of artificial polyadenylation (Isakova et al., 2021; Wulf et al., 2022). However, only RNAs that contain a 3’OH can be artificially polyadenylated, preventing the detection of RNAs with blocked 3’. TGIRT-seq (Mohr et al., 2013; Xu et al., 2019) leverages a group II intron reverse transcriptase to synthesize full length cDNAs, including tRNAs (Zheng et al., 2015), without the need for 3’ adapter ligation. TGIRT-seq detects both small and long RNAs, but suffers from sequence bias, as well as from strong adapter dimers and from the formation of cDNA concatamers (Xu et al., 2019). The recently developed mim-tRNAseq protocol significantly improved TGIRT-mediated cloning of tRNAs, which are known to be heavily modified both internally and at the 3’ (Suzuki, 2021), but at the expense of chemical deacylation of 3’ tRNA ends (Behrens et al., 2021), which could cause unwanted RNA fragmentation. OTTR-seq (Up-ton et al., 2021) utilizes the “template jumping” activity of the truncated *B. mori* R2 RT to synthesize full length cDNA with 3’ and 5’ adapter sequences. This improved the detection of several RNA classes, including tRNAs (Gustafsson et al., 2022), but it is unclear whether OTTR-seq can efficiently clone RNAs larger than 200 nts. In addition, the terminal nucleotidyl transferase activity of the *B. mori* R2 RT towards the template RNA, necessary for primer duplex hybridization (Upton et al., 2021), requires free (3’OH) ends. Furthermore, the *B. mori* R2 RT is not yet commercially available, making its integration into sequencing protocols more difficult.

LIDAR captured a synthetic RNA with a biotinylated 3’ with comparable efficiency as a control RNA with a 3’OH, whereas the ligation-based method completely missed the blocked RNA (**Fig. 4B**). At the same time, LIDAR libraries from total RNA contained inserts from a variety of small and long RNAs, including full-length tRNAs and tDRs that could not be ligated, likely due to the presence of an amino acid in 3’ (see below). Thus, LIDAR is an effective, truly ligation-independent alternative to the methods cited above, with the added advantage of its simplicity and the commercial availability of all required reagents.

### A new class of 3’ tDRs?

Several regulatory functions have been assigned to tDRs (Su et al., 2020). For example, 5’ tiRNAs generated by angiogenin cleavage sustain cell proliferation, at least in breast cancer cells (Honda et al., 2015) and, in plants, 3’ tDRs participate in retrotransposon silencing (Schorn et al., 2017). tDRs found in sperm have been implicated in the epigenetic inheritance of metabolic disorders (Chen et al., 2016; Sharma et al., 2016). In these reports, ligation-based methods showed predominance of 5’ tDRs in mature sperm and very few 3’ tDRs. However, the presence of 3’ tDRs in sperm was detected by northern blot (Sharma et al., 2018), the recent OTTR-seq method (Gustafsson et al., 2022), and, now, LIDAR (**Fig. 5C**). Attempts to recapitulate the epigenetic transmission of metabolic information using a synthetic 5’ tDRs based on those detected in sperm by a ligation-based method proved to be unsuccessful (Chen et al., 2016), indicating that our knowledge of sperm RNA content may, in fact, be far from complete.

LIDAR detected the presence of blocked 3’ tDRs in both ESC and sperm, which we speculate to be aminoacylated, although only one example has been reported so far (Liu et al., 2021). It is also possible that some of the 3’ tDRs detected by LIDAR may be instead part of nicked mature tRNA, as proposed in some recent studies (Chen and Wolin, 2023; Costa et al., 2023). Regardless of their origin, we believe that loaded 3’ tDRs may represent an overlooked class of regulatory RNAs with potentially important and yet unknown regulatory functions, which can now be more thoroughly investigated thanks to the introduction of LIDAR.

Many reads in LIDAR libraries from various sources, including sperm, mapped to mitochondrial tRNAs **(Fig. 1F, 5A**). Their persistence after enrichment for blocked RNA species (**Fig. S4E, 5B**) suggests that they are charged with amino acids, explaining why ligation-based methods failed to capture them. Non-canonical functions of mitochondrial tDRs have been proposed (Shaukat et al., 2021), but their role is still unclear, especially in the context of sperm maturation/ function, and in need of further investigation.

### Future improvements and alternatives

In its current form, LIDAR cannot be used for single cell analysis due to the substantial amount of adapter dimer contaminations when starting from less than 10 ng of RNA. Post-library depletion methods, such as DASH (Dynerman et al., 2020; Gu et al., 2016), might be used to remove adapter dimers, and therefore increase sensitivity of LIDAR to the point of allowing its application in single cells. Along similar lines, a large proportion of LIDAR reads mapped to rRNAs. If more sensitivity towards non-rRNA species is required, rRNA depletion methods (Herbert et al., 2018) could be easily integrated into the LIDAR protocol.

Despite several optimization attempts, some miRNAs that could be detected via ligation were missed by LIDAR. One possibility is that miRNA end modifications (5’P and 3’OH) (Crocker et al., 2022) represent ideal substrates for ligation, while template switching is more favorable on 5’ capped or 5’OH RNAs (Wulf et al., 2019). In cases when detection of the full spectrum of 5’P RNAs is crucial, RNA could be chemically capped (Wulf et al., 2022) or dephosphorylated before LIDAR library construction. One type of RNA that might be missed by LIDAR are circular RNAs (Yang et al., 2022), since the lack of a linear 5’ end likely favor rolling-circle cDNA synthesis over template switching (Das et al., 2019). To enhance circular RNA detection with LIDAR, it should be possible to enrich circular RNAs and then perform gentle hydrolysis, as recently described (Rahimi et al., 2021).

In addition to 3’ tDRs, LIDAR also revealed the presence of blocked 5’ tDRs (**Fig. S5E**). Because, although unlikely, internal priming events on full-length tRNAs generating 5’-tDRs-like reads cannot be formally excluded, we chose to focus most of the analyses on the 3’ tDRs. If a precise and sensitive cloning protocol for 5’ tDRs is desired, a combination of TGIRT-mediated 3’ (Xu et al., 2019) and Smart-mediated 5’ (Hagemann-Jensen et al., 2020) template switches may be the solution. Adding a tagmentation step may also increase coverage of longer RNAs.

### Conclusions and outlook

LIDAR is a powerful, simple, and fast RNA-seq method that allows comprehensive characterization of coding and noncoding transcriptomes from limiting amounts of materials with commercially available reagents. LIDAR allowed us to capture, with a simple and fast protocol, a large variety of RNAs in mouse ESC, NPC, and sperm, including full-length tRNAs and 3’ tDRs with blocked 3’ termini that previous methods failed to detect. While the functional implications of the presence of these 3’-blocked tDRs in sperm are currently unknown, we noticed that they are of the same size as the RNA population responsible for the epigenetic transmission of a metabolic disorder through the mouse germline (Chen et al., 2016; Sharma et al., 2016).

Full-length tRNAs and tDRs might not be the only RNAs with blocked 3’ ends. Our data suggest the intriguing possibility that 3’ blocked small RNAs might also be formed from other classes of transcripts, such as snoRNAs and rRNAs (**Fig. 4G, S5C**). Given that LIDAR has the potential to capture RNAs with any type of 3’ modifications, known or unknown, without the need to develop dedicated chemical strategies for their removal, we believe it will become an essential tool to investigate these yet unexplored RNAs.

## Materials and Methods

### Synthetic RNAs

The synthetic 20 nts and 50 nts fragments were synthesized by Integrated DNA technologies (IDT) (**Table S1**). For 3’ biotinylated RNAs, we performed a purification step using Dynabeads MyOne Streptavidin C1 beads (Invitrogen, Cat. No 65001) followed by TriPure (Roche, Cat. No 11667165001) purification to ensure all RNAs used in the reaction contained biotin.

### Cell culture and NPC differentiation

C57BL/6 mouse ESC were purchased from ATCC (SCRC1002) and maintained onto 1% gelatin-coated (Sigma-Aldrich, Cat. No G1393) dishes in DMEM knockout medium (Life Technologies, Cat. No 10829-018) supplemented with 15% ES-grade fetal bovine serum (Gibco, Cat. No 16141079), 1% GlutaMAX (Thermo Scientific, Cat. No 35050061), 1% non-essential amino acids (Sigma-Aldrich, Cat. No M7145), 0.5% penicillin/streptomycin solution (Sigma-Aldrich, Cat. No P0781), 110 μM β-mercaptoethanol (Gibco, Cat. No 21985023), 100 units/mL leukemia inhibitory factor (LIF) (Millipore Sigma, Cat. No ESG1107), and 2i [3 μM GSK-3 inhibitor XVI (Cat. No 361559), 1 μM MEK1/2 Inhibitor III (Cat. No 444966)].

Prior to NPC differentiation, ESC were adapted for > 2 weeks to N2B27 serum-free medium consisting of 1:1 mix of neurobasal medium (Gibco, Cat. No 1103049) and DMEM/F12 (Gibco, Cat. No 11320033), supplemented with 1X N2 (Gibco, Cat. No 17502048), 1X B27 (Gibco, Cat. No 17504044), 1% GlutaMAX, 1% non-essential amino acids, 0.5% penicillin/streptomycin, 55 μM β-mercaptoethanol, 100 units/mL LIF, and 2i. NPC differentiation was performed as previously described (Petracovici and Bonasio, 2021). Briefly, EpiLCs were induced in N2B27 serum-free medium without 2i and LIF and with 40 μg/mL of BSA and 10 ng/ mL bFGF (R&D Systems, Cat. No 3139-FB-025) for 72 h. EpiLCs were then treated with 500 nM SAG (Calbiochem, Cat. No 566661) for 48 h to generate NPC.

### Sperm collection and isolation

Cauda epididymides were dissected from adult (8-12 weeks old) male FVB/NJ mice into a dish with 1.5 ml prewarmed Whitten’s media. Caudal fluid was gently squeezed from the tissue and left to incubate in the dish for 10 mins at 37°C. After incubation, sperm containing media was transferred to a 1.5 ml tube and sperm was allowed to ‘swim up’ for 10 min at 37°C. The sperm containing media was collected, leaving the bottom 50 ml and sperm was pelleted by centrifugation (10,000 x g, 5 min, 4°C). The sperm pellet was washed with 1 x PBS and somatic cells were removed by incubation in somatic cell lysis buffer (0.01% SDS and 0.005% Triton-X) for 10 min on ice. After a final wash in PBS, the sperm pellet was snap frozen in liquid nitrogen and stored at -80°C.

### RNA extraction and fractionation

For ESC and NPC, cell pellets were resuspended in 1 mL of TriPure and RNA extracted following standard protocol. DNA was digested with Turbo DNase-I (Invitrogen, Cat. No AM2238) at 37°C for 30 min, RNA was re-purified using TriPure, and resuspended in modBTE (10mM Bis/Tris pH 6.7, 0.1 mM EDTA). For mouse sperm, total RNA was extracted using the same method previously described for epididymosomes (Conine et al., 2018). Briefly, sperm were resuspended in 120 µL of water and 66 μL of sperm lysis_buffer were added [120mM Tris-HCl (pH 8), 6.4 M Guanidine-HCl, 5% Tween-20, 5% Triton-X-100, 120 mM EDTA]. Proteins were digested by adding 6.6 μL of 20 mg/mL proteinase K and 6.6 μL of 1M DTT, followed by incubation at 60°C for 15 min under 600 rpm constant shaking. Volume was adjusted with water to 400 μL and 400 μL of TriPure were added. RNA was extracted by adding 120 μL of BCP phase separation reagent (Molecular Research Center, Cat. No BP151) and precipitated with isopropanol. DNA was digested with Turbo DNase-I at 37°C for 30 min, RNA was re-purified using TriPure, and resuspended in modBTE .

For enrichment of RNAs < 200 nts, the Zymo RNA Clean and Concentrator-5 kit (Cat. No.) was used, starting from 1 µg of total RNA. For enrichment of RNAs < 50 nts, 18 µg of total RNA were run on a denaturing 12% polyacrylamide-urea gel, and the section between 10 nts and 50 nts markers was excised. Gel pieces were shredded trough pierced 0.5 mL tubes, and RNA was eluted from the gel by overnight incubation in 400 μL RNA elution buffer (10mM Bis/Tris pH 6.7, 300 mM NaCl, 10 mM EDTA) at 4°C with constant rotation. Eluate was filtered through 5 μm PVDF spin filters (EMD Millipore, Cat. No UFC30SV00), and RNA was precipitated by adding 1 μL glycoblue (Invitrogen, Cat. No AM9516), 40 μL 3 M NaOAc pH 5.2, and 1.1 mL of icecold 100% EtOH, followed by incubation at -80°C for 1 h. RNA was pelleted at 20,000 g for 30 min at 4°C, washed once with 1 mL of 70% EtOH, once with 1 mL of 80% EtOH, air-dried for 5 min, and resuspended in modBTE buffer.

### LIDAR and 3’-LIDAR library preparation

The desired amount of total and fractionated RNAs was diluted to 1 μL and mixed with 0.4 μL of 10 μM LIDAR_RT_ primer (see **Table S1** for all oligonucleotide sequences). To anneal the LIDAR_RT_primer to the template RNA, samples were heated to 65°C for 5 min, then cooled to 4°C (0.5°C/s). To initiate RT, 6.28 μL of RT_mix [25mM Tris-HCl pH 8.3, 20 mM NaCl, 2.5 mM MgCl_2_, 8 mM DTT, 5% PEG-8000, 0.5 mM dNTPs, 1 mM GTP, 0.5 U/μL murine RNase inhibitor (New England Biolabs, Cat. No M0314S), and 2U/μL Maxima H-minus RT (Thermo Scientific, Cat. No EP0752); concentrations refer to a final volume of 8 μL] were added and samples were incubated at 25°C for 10 min. To further promote cDNA synthesis and template switch, temperature was raised to 42°C, 0.32 μL of 50 μM LIDAR_TSO_mix (Table S1) were added, and samples were incubated at 42°C for 80 min, followed by 10 cycles at 50°C for 2 min and 42°C for 2 min, then at 85°C for 5 min. To pre-amplify and add adapters to cDNA, 12 μL of KAPA_mix [1X KAPA HiFi HotStart buffer Ready Mix (Roche, Cat. No KK2601), 0.5 μM LIDAR_preamp_f, and 0.1 μM LIDAR_preamp_r; concentrations refer to a final volume of 20 μL] were added and samples were incubated with the following PCR cycling conditions: denaturation (95°C for 3 min), 18 x (98°C for 20 s, 70°C for 30 s, 72°C for 30 s), final extension (72°C for 5 min). Quality of adapter-containing libraries was assessed by loading 5 uL on a 2% agarose gel. To generate the final libraries, 2 μL of pre-amplified were diluted to 37 μL with 10mM Tris-HCl pH 8, 0.5 μL of each 10 μM custom Nextera indexing primers was added, followed by 12 μL of Q5_mix [1X Q5 buffer, 0.5 mM dNTPs, 0.02 U/µL Q5 High-Fidelity DNA Polymerase (New England Biolabs, Cat. No M0491); concentrations refer to a final volume of 50 μL]. Samples were incubated with the following PCR cycling conditions: denaturation (98°C for 30 sec), 18 x (98°C for 10 s, 65°C for 20 s, 72°C for 20 s), final extension (72°C for 2 min). Indexed libraries were purified using 2.3X SPRI beads (Beckman Coulter, Cat. No B23319) and eluted in 50 μL of TE buffer (10mM Tris-HCl pH 8, 1 mM EDTA).

For 3’-LIDAR, the LIDAR-3_RT_oligo was generated by mixing equimolar amounts of LIDAR_RT_primer **Table S1**) and LIDAR-3_RT_antisense (**Table S1**) followed by heating at 95°C for 5 min, and a slow cool down to 25°C (0.1°C/s). To prepare 3’-LIDAR libraries, 1 μL of input RNA was first denatured at 70°C for 2 min, then temperature was lowered to 50°C and 0.4 μL of 1 μM LIDAR-3_RT_oligo were added. Samples were incubated at 50°C for further 2 min, then temperature was lowered to 4°C (0.5°C/s). To initiate RT, 6.28 μL of RT_mix were added and samples were incubated at 25°C for 10 min. To further promote cDNA synthesis and template switch, temperature was raised to 42°C, 0.32 μL of 50 μM LIDAR_TSO_mix were added, and samples were incubated at 50°C for 80 min, followed by 10 cycles at 55°C for 2 min and 50°C for 2 min, then at 85°C for 5 min. LIDAR-3_RT_antisense was digested by adding 1 μL of USER II enzyme (New England Biolabs, Cat. No M5508S) and incubating at 37°C for 30 min. USER II was heat inactivated by incubation at 65°C for 10 min. To pre-amplify and add adapters to cDNA, 12 μL of KAPA_mix (1X KAPA HiFi HotStart buffer ready mix, 2 μM LIDAR_preamp_f, and 0.5 μM LIDAR_preamp_r; concentrations refer to a final volume of 20 μL) were added and samples were incubated with the following PCR cycling conditions: denaturation (95°C for 3 min), 6x (98°C for 20 s, 63°C for 30 s, 72°C for 30 s), 12x (98°C for 20 s, 72°C for 50 s), final extension (72°C for 5 min). Quality of adapter-containing libraries was assessed by loading 5 µL on a 2% agarose gel. To generate the final libraries, 2 μL of pre-amplified libraries were diluted to 37 μL with 10 mM Tris-HCl pH 8, 1 μL of each 10 μM custom Nextera indexing primers was added, followed by 12 μL of Q5_mix (1X Q5 buffer, 0.5 mM dNTPs, 0.02 U/ µL Q5 high-fidelity DNA polymerase; concentrations refer to a final volume of 50 μL]. Samples were incubated with the following PCR cycling conditions: denaturation (98°C for 30 sec), 7x (98°C for 10 s, 65°C for 20 s, 72°C for 20 s), final extension (72°C for 5 min). Indexed libraries were purified using 2.4X SPRI beads and eluted in 50 μL of TE buffer (10 mM Tris-HCl pH 8, 1 mM EDTA).

LIDAR and 3’-LIDAR libraries were analyzed on a 2% agarose gel and quantified using NEBNext Library Quant Kit for Illumina (New England Biolabs, Cat. No E7630L). Libraries were sequenced on an Illumina NextSeq500.

### Small RNA-seq and mRNA-seq library preparation

Small RNA libraries were prepared using the NEBnext small RNA library kit for Illumina (New England Biolabs, Cat. No E7330S), following the standard protocol with the following parameters: 1) 1:2 dilution of 3’SR Adaptor, SR RT Primer, and 5’SR Adaptor, 2) 15 indexing PCR cycles, 3) cleanup of final libraries with QIAgen MinElute PCR purification kit (QIAgen, Cat. No 28004), followed by a second cleanup with 2X SPRI. Libraries were analyzed on 6% non-denaturing polyacrylamide gels.

For mRNA-seq libraries, polyA RNA was enriched from 2 μg total RNA using Dynabeads Oligo(dT)25 (Invitrogen, Cat. No 61002). polyA-enriched fraction was used for library construction using the NEBnext Ultra II Directional Library Prep Kit for Illumina (New England Biolabs, Cat. No. E7760L), following the standard protocol but using half the reaction volumes recommended. Libraries were analyzed on 2% agarose gels.

Both small RNA-seq and mRNA-seq libraries were quantified using NEBNext Library Quant Kit for Illumina (New England Biolabs, Cat. No E7630L). Libraries were sequenced on an Illumina NextSeq500.

### Enrichment of blocked RNAs

To enrich RNA species with blocked 3’, input RNA (300 ng of total ESC RNA, 75 ng of < 200 nts ESC RNA, or 75 ng of total sperm RNA) was first diluted in 38.5 μL of water and heat-denatured at 70°C for 2 min, followed by quick cool down at 4°C. To generate 5’P and 3’OH ends, 11.5 μL of PNK mix [1X T4 PNK reaction buffer, 1 mM ATP, 1 U/μL T4 PNK (New England Biolabs, Cat no M0201S); concentrations refer to a final volume of 50 μL] were added, and samples were incubated at 37°C for 30 min. 1 mL of TriPure was added, RNA was purified following the standard protocol, and resuspended in 14.5 µL of water. To polyadenylate the end-repaired RNA, 5.5 μL of EPAP mix [1X *E. coli* Poly(A) Polymerase Reaction Buffer, 1 mM ATP, 0.25 U/μl *E. coli* Poly(A) Polymerase (Cat no, NEB), 2 U/μl RNase inhibitor, murine (Cat no, NEB); concentrations refer to a final volume of 20 μL] were added, and samples were incubated at 37°C for 10 min. Polyadenylation reaction was stopped by adding EDTA to a final concentration of 10 mM. To deplete polyadenylated RNAs, 100 μL of Dynabeads Oligo(dT)25 were added to the sample and hybridization was carried at 25°C for 25 min. Flow-through was collected and RNA was purified with TriPure following standard protocol. As control, bead-bound RNA was also extracted using the same procedure. Fractionated RNA was analyzed on 9% denaturing (urea) polyacrylamide gels.

### Data processing

#### mRNA-seq

Adapters were trimmed using TrimGalore with default parameters (ver 0.6.4_dev using Cutadapt version 4.2, https://github.com/FelixKrueger/TrimGalore and DOI:10.14806/ ej.17.1.200), retaining reads with a minimum length of 15 bp for both R1 and R2.

#### NEB

Adapters were trimmed using TrimGalore, retaining paired reads with a minimum length for both R1 and R2 of 5 bp, and R1 singletons with a minimum length of 20 bp (*trim_ galore –length 5 --paired --retain_unpaired -r 20 -r2 100*). The 3’ R2 adapter GATCGTCGG was further trimmed using cutadapt (*cutadapt -A GATCGTCGG --minimum-length 5 --pair-filter=any* or *cutadapt -a GATCGTCGG --minimum-length 5*). Paired reads that overlapped by at least 8 bp were connected using COPE (ver 1.2.5) (Liu et al., 2012) in simple connect mode (*cope -s 33 -m 0 -l 8*). The connection of read pairs at this step results in some paired reads and some single reads, which were processed in parallel during mapping.

#### LIDAR

Reads were processed with TrimGalore and COPE as above, excluding the 3’ R2 adapter trimming. UMIs were extracted using umi_tools extract (ver 1.1.2). To account for the variable length UMIs, three X bases were added to the beginning of each read, then UMIs were extracted with the regex “.*(?<UMI_1>.{7})CG(?P<UMI_2>.{2})AG(?P<Umi_3>.{2})GGG”, producing the following patterns for the 4 variable length options, where numbers refer to the six UMI bases:

+0 XXX TG 12 CG 34 AG 56 GGG - insert -> XXXTG123456-

insert

+1 XXX VHG 12 CG 34AG 56 GGG -insert -> XXVHG123456-

insert

+2 XXX VATG 12 CG 34 AG 56 GGG - insert ->

XVATG123456-insert

+3 XXX VCMTG 12 CG 34 AG 56 GGG - insert ->

VCMTG123456-insert

In this way, 151,552 UMIs can be encoded, taking into account the 4 different variable length options, the 0–3 bases preceding the first UMI bases, and the 6 UMI bases.

After UMI extraction, the remaining constant bases (CGAGGGG) were trimmed from the read using cutadapt (*cutadapt --minimum-length 5 -g CGAGGGG* for single reads or *cutadapt --minimum-length 5 -g CGAGGGG --pair-filter=any* for paired reads), followed by another trimming step to remove any reads with an occurrence of the TSO or its reverse complement, to stringently remove reads resulting from adapter dimers (*cutadapt -e 3 -b CGTCAGATGTG- TATAAGAGACAG --discard-trimmed*).

### Synthetic RNAs

For samples containing only synthetic RNAs of known sequence, reads were processed as above for LIDAR and NEB, but with a length cutoff of 15 bp required for R1 and R2 during adapter trimming (*trim_galore --paired --length 15*). Only reads collapsed with COPE were retained, as R1 and R2 for the 20-nt or 50-nt synthetic RNAs should overlap. LIDAR samples were processed to extract UMIs, as described in *Read processing* section.

The 8N mix (**Fig. 1D–E, S1C–D**) contains a mixture of 20 nts and 50 nts synthetic RNA oligos, with the first 12 or 42 bp constant and a random 8 bp following. Reads capturing the 20 nts RNA were retained if they contained the constant first 12 bp with up to one mismatch and had a total length ≤ 20 bp, while reads capturing the 50 nts RNA were retained if they contained the constant first 42 bp with up to three mismatches and had a total length ≤ 50 bp. RPMs were calculated from all reads that passed the adapter trimming step. SeqLogos were generated for the 8N bases by calculating a position weight matrix for all reads with a base at the positions indicated (**Fig. S1D**).

The 3’OH/3’ blocked oligos mix (**Fig 4B, S4A**) contains a mixture of 20 nts and 50 nts synthetic RNAs, some with a 3’OH and others with a blocked 3’ end. After read processing, reads were aligned to a reference composed of the oligo sequences using bowtie2 (ver 2.5.0) (Langmead and Salzberg, 2012) with default parameters, and LIDAR samples were deduplicated. Only reads with an insert length of ≥ 15 bp were considered. RPMs were calculated from all reads that passed the adapter trimming step.

For the LIDAR/3’-LIDAR comparison (**Fig. S3C**), all reads with a fragment length ≥ 15 bp mapping to the 3’OH oligos were considered. The plot represents the % of these reads that end at the 3’ end of the oligo.

### Mapping (mRNA-seq, NEB, LIDAR)

Reads were mapped in multiple passes to sequentially identify rRNA reads and mapping to ncRNA before mapping to the main genome. Reads mapping at each step were removed before the next step. Only reads with an insert length ≥ 15 bp were retained for downstream analysis.

Locations in the genome predicted to be rRNA repeats were identified using RepeatMasker tracks downloaded from the NCBI Table Browser (group=all talbes, table=rmsk, repClass=rRNA) and masked in the main genome, and a consensus scaffold of rRNA repeats (BK000964.3) was added as a separate scaffold. Genomic loci corresponding to snoRNA and snRNA genes in the GENCODE annotation (ver M27) (Frankish et al., 2019), as well as piRNA loci predicted by piRbase (release v3.0 gold standard set) (Wang et al., 2022) and tRNA from GtRNAdb and tRNAscan-SE, as curated in Behrens et al. (Behrens et al., 2021; Chan and Lowe, 2016; Lowe and Chan, 2016), were also identified. Locations in the genome corresponding to rRNA and tRNA were masked in the main genome and their sequences were added as separate scaffolds.

1. Reads were mapped with STAR (ver 2.7.10a_alpha_220601) (Dobin et al., 2013): to a consensus scaffold of rRNA repeats (BK000964.3) with the following parameters: --outFilterMatchNmin 16 --alignIntronMax 1 –outFilterScoreMinOverLread 0.9 --outFilterMatchNminOverLread 0.9 --outFilterMismatchNoverLmax 0.05.
2. Remaining reads were mapped with bowtie2 (ver 2.5.0) to snoRNA sequences described above in --very-sensitive mode. Paired reads mapping in proper pairs with insert size > 15 bp and single reads with insert size > 15 bp were retained as mapping.
3. Remaining reads were mapped as in step 2 to snRNA.
4. Remaining reads were mapped as in step 2 to piRNA.
5. Remaining reads were mapped to tRNA using gsnap (ver 2019-02-26) (Wu and Nacu, 2010)2010, in SNP-tolerant alignment mode with parameters and databases described in in Behrens et al. (Behrens and Nedialkova, 2022; Behrens et al., 2021), including pre-built references of tRNA predictions from GtRNAdb and modifications from MODOMICS (Boccaletto et al., 2018). The following parameters were used: -D <TRNA genome directory> -d <TRNA genome> -V <TRNA genome index> -v <TRNA modification database> --ignore-trim-in-filtering 1 --format sam --genome-unk-mismatch 0 unmapped --md-lowercase-snp --max-mismatches 0.075. Reads mapped concordantly, uniquely or multimapped, were retained and all others were considered unmapped.
6. Mapping of remaining paired and single reads

a. Remaining paired reads were mapped to the modified GRCm39 genome, as described above, using STAR with the following parameters: --peOverlapNbasesMin 5 --alignIntronMax 20 –alignIntronMax 100000 –outFilterMismatchNoverLmax 0.05 --outFilterScoreMinOverLread 0.9 --outFilterMatchNminOverLread 0.9 --out- FilterMatchNmin 16. Unmapped reads or reads with insert length smaller than 100 were mapped to the modified GRCm39 genome using bowtie2 in --very-sensitive mode.
b. Remaining single reads were mapped to the modified GRCm39 genome with bowtie2 in --very-sensitive mode.
7. All mapped paired reads (rRNA, snoRNA, snRNA, piRNA, tRNA, main genome) were merged, and all mapped single reads were merged.
8. LIDAR samples were deduplicated using umi_tools dedup with --method unique (ver 1.1.2).
9. Only reads with fragment length ≥ 15 bp were included in downstream analyses.

### Mispriming analysis

The percentage of reads with a mispriming event (**Fig. S1G**) was defined as the percentage of reads with the UMI sequence (tgNNcgNNagNNGGG) templated in the DNA, indicating a potential TSO strand invasion (Hahaut et al., 2022). Bases 15 nt upstream of the read start were considered, with 0–2 allowed mismatches.

### Read counting for genes and biotypes

A custom annotation was created using RefSeq Annotation Release 109 (GRCm39) with miRNA removed, and miRNA from miRbase (v22) (Kozomara et al., 2019) added; only mature miRNA were included, and coordinates were converted from GRCm38 to GRCm39 using the LiftOver tool (Hinrichs et al., 2006). Reads mapping to this annotation were counted using an in-house script based on the GenomicRanges (ver 1.50.2) (Lawrence et al., 2013) function SummarizeOverlaps, counting the number of reads overlapping exons of genes with counting mode IntersectionNotEmpty. The command used to compute counts was *assay(summarizeOverlaps(annotation,bam_file, ignore. strand = F, singleEnd = T, param = scanBamParam(flag = scanBamFlag(isSecondaryAlignment = FALSE)), mode = ”IntersectionNotEmpty”))*, with singleEnd = F for paired-end reads. Any reads mapping to the ncRNA (snoRNA, snRNA, piRNA, tRNA, rRNA) scaffolds as described above were counted towards the total for those genes. TPMs were calculated to take library size into consideration.

To determine the % of reads in each sample mapping to each biotype (**Fig. 1F, 5A, S1H, S1J, S2A, S4E**), all genes for each biotype were collapsed, alleviating issues where multiple genes of one biotype overlapped each other, making read assignment to the specific gene impossible. Any regions of the genome that contained genes of two different biotypes were marked as “ambiguous coding” or “ambiguous non-coding” depending on the presence of a protein-coding gene in that region. Reads mapping to the consensus rRNA scaffold BK000964.3 were ignored for these analyses. The biotypes considered “other” were pseudogene, transcribed pseudogene, misc_RNA, guide RNA, antisense RNA, RNase P RNA, telomerase RNA, RNase MRP RNA, V segment, V segment pseudogene, D segment, J segment, C region, J segment pseudogene, Y RNA, scRNA, and ncRNA pseudogene. Reads mapping to each biotypes were counted as above.

The number of snoRNA and miRNA detected per million reads (**Fig. S1K**) was determined by randomly selecting 1 million reads that map to annotated genes for each sample, with proportions reflecting the underlying read count tables, then counting the number of snoRNA or miRNA with at least one read. The number of genes detected in length bins (**Fig. S2B**) was determined by first generating lists of expressed genes in each length bin, considering any gene with an average TPM ≥ 1 among all samples (LIDAR and ligation libraries; < 50 nt, < 200 nt, and total fractions, *n* = 3 for each condition). The % of expressed genes in each length bin was then calculated, with a threshold of TPM ≥ 1 for detection.

### Differential expression

Differential expression between ESC and NPC (**Fig. 2B**) was performed using DESeq2 (ver 1.38.3) (Love et al., 2014). Genes with an adjusted *p* value < 0.05 were considered differentially expressed.

Differential expression for tDRs (**Fig. 4H**, **5E**) was performed only considering reads mapped to the indicated tDRs, generating results that indicate differences in frequency of genes between samples.

### tRNA analysis

#### Classification of tRNA reads into full-length/fragment types

For each tRNA considered in the analysis, the position of each canonical tRNA base, with numbering as provided in Sprinzl et al. (Sprinzl and Vassilenko, 2005) was annotated using a multiple species alignment of each tRNA to identify canonical bases. Non-templated CCA bases were included in the reference.

Reads mapping to tRNA were classified as full-length, 5’ tRF, 5’ tiRF, internal tRF (i-tRF), 5’ tRF, 3’ tRF, or “other” reads using the following definitions:

1. Full-length tRNA read: read spans from position 1 at the 5’ end to within 5 bp of the 3’ end (including the CCA) of the tRNA.
2. 5’ tRF: read spans from position 1 at the 5’ end to ≤ canonical base 31
3. 5’ tiRF: read spans from position 1 at the 5’ end to ≤ canonical base 35, excluding reads classified as 5’ tRFs
4. i-tRF: read represents an internal fragment, spanning from ≥ 10 bp from 5’ end and ≥ 10 bp from 3’ end of tRNA
5. 3’ tRF: read spans from ≥ canonical base 48 to within 5 bp from the 3’ end of the tRNA
6. 3’ tiRF: read spans from ≥ canonical base 35 to within 5 bp from the 3’ end of the tRNA, excluding reads classified as 3’ tRFs

#### Misincorporation analysis (**Fig. 3E, 3F, S4D, S5F**)

For each cytosolic tRNA, the percentage of reads with a base differing from the reference or known SNP database was computed for each canonical position, as described above. Only tRNAs with at least 5 reads assigned in all replicates of at least one condition (for example, LIDAR 200 nt ESC) were included in the heatmaps; positions without coverage were considered NA and not included in mean values plotted on the heatmap.

Start and end position of fragments **(Fig. 3C, S3G, S5E)**

All reads considered 5’ fragments (reads spanning from position 1 at the 5’ end of tRNA to ≤ canonical base 35) and 3’ fragments (reads spanning from ≥ canonical base 48 to within 5 bp from the 3’ end of tRNA) were considered. The end position in terms of tRNA canonical positions was calculated for 5’ fragments, while the start position was calculated for 3’ fragments.

#### tRNA coverage by codon (**Fig. 4D**)

For each tRNA, the coverage for each canonical position, as described above, was calculated. Coverage over the variable region was collapsed by taking the mean of all variable positions. In the case of a tRNA missing canonical bases, coverage was calculated as the mean of the closest present canonical bases to the left and right. Coverage was then collapsed by anticodon by taking the sum of all tRNAs with each anticodon and converted to RPMs based on the total number of reads mapped in each sample.

#### Data visualization

All computational plots were generated using ggplot2 (ver 3.4.2, H. Wickham. ggplot2: Elegant Graphics for Data Analysis. Springer-Verlag New York, 2016) with viridis (ver 0.6.2, Simon Garnier, Noam Ross, Robert Rudis, Antônio P. Camargo, Marco Sciaini, and Cédric Scherer (2021). Rvision Colorblind-Friendly Color Maps for R) or scico (ver 1.3.1, Pedersen T, Crameri F (2022). scico: Colour Palettes Based on the Scientific Colour-Maps) color palettes, except heatmaps that were visualized using pheatmap (ver 1.0.12, Kolde R (2019) https://CRAN.R-project.org/pack-age=pheatmap). SeqLogos were created using the SeqLogo package (ver 1.62.0, Bembom O, Ivanek R (2022).

## Data Availability

All sequencing data generated in this manuscript have been deposited in NCBI GEO (accession number: GSE233343).

## Acknowledgments

The authors thank Robert Warneford-Thomson for technical support, and the Bonasio lab for constant feedback and for comments on the manuscript. R.B. acknowledges support from the NIH (R01GM127408; R01GM138788). N.A.T. is supported by the Lalor Foundation Postdoctoral Fellowship. J.E.W. is supported by National Institutes of Health grant R35-GM119735 and Cancer Prevention & Research Institute of Texas grant RR210031. J.E.W. is a CPRIT Scholar in Cancer Research. C.C.C. is supported by the Pew Biomedical Scholars Award.

## Author Contributions

Conceptualization: A.S. and R.B.; methodology: A.S., E.J.S., and R.B.; formal analyses: E.J.S; investigation: A.S., N.A.T., and J.E.W.; data curation: E.J.S.; writing – original draft: A.S. and R.B.; writing – review & editing: A.S., E.J.S., N.A.T., C.C.C., J.E.W., and R.B.; visualization: A.S., E.J.S., and R.B.; supervision: R.B., and C.C.C.; project administration: R.B.; funding acquisition: R.B., J.E.W., and C.C.C.

---

**Figure S1.**
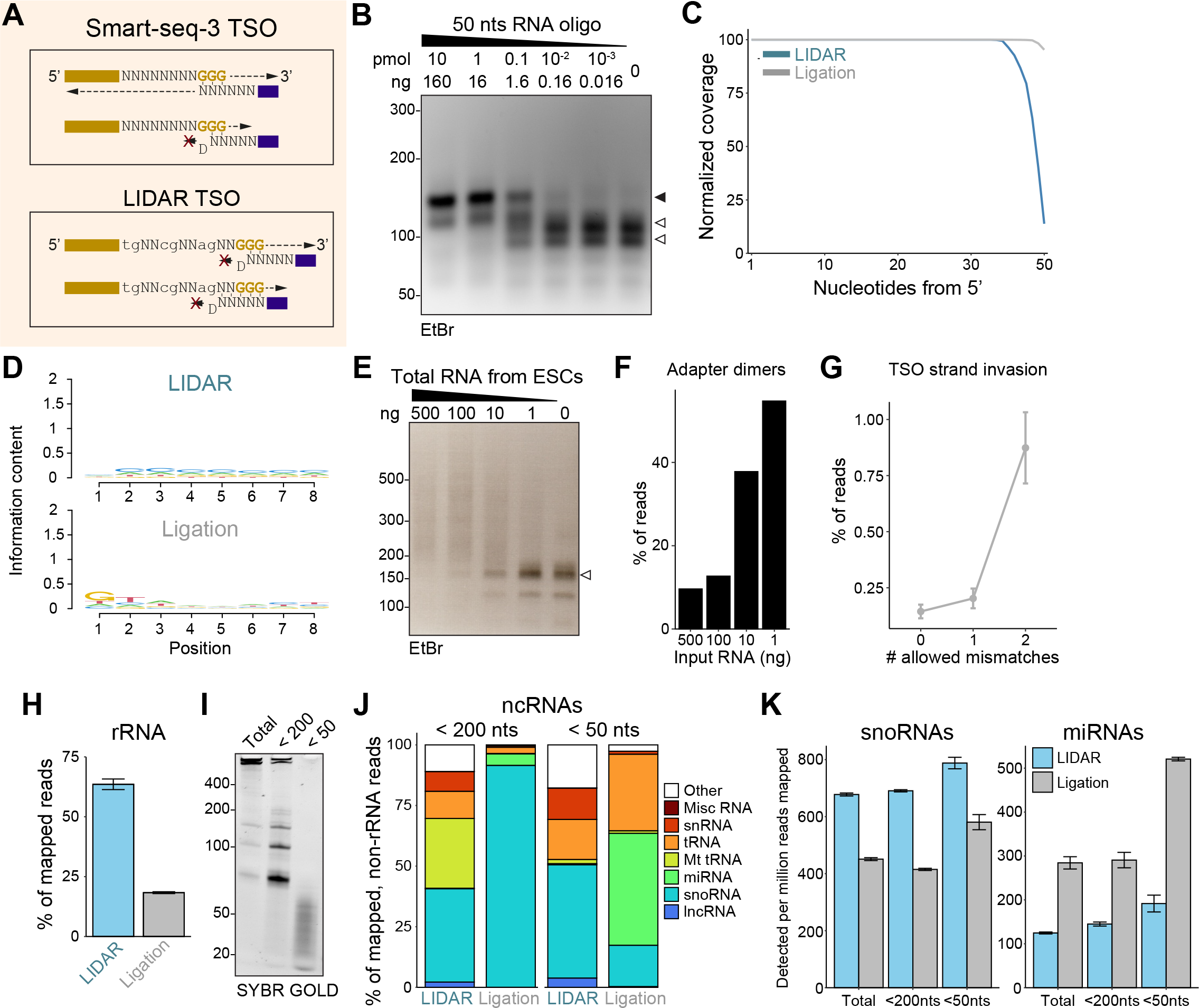
**Additional comparisons of LIDAR and ligation-based libraries** (A) Scheme of random and quasi-random priming in the presence of the Smart-seq3 (top panel) or the LIDAR TSO (bottom panel). The presence of a D nucleotide (A, G, or T) at the 3’ of the quasi-random hexamers prevents RT priming on both TSOs. (B) Agarose gel electrophoresis of pre-amplified (step III of Fig. 1A), LIDAR libraries constructed from the indicated amounts of a 50 nts RNA oligo. Black arrowhead indicates productive libraries; white arrowheads indicate adapter dimers. (C) Read coverage of a 50 nts RNA oligo cloned with LIDAR (blue) or with a conventional ligation-dependent protocol (gray). Read density is expressed as % of coverage across the constant region. (D) Information content of sequence composition of the last 8 nts of a 20 nts RNA cloned with LIDAR or ligation-dependent protocol. Numbers indicate base position from the constant portion of the oligo to the 3’ end (position 8 is the terminal 3’ nucleotide). (E) Agarose gel electrophoresis of final (step IV of Fig. 1A) LIDAR libraries starting from decreasing amounts of total RNA from ESC. White arrowhead indicates adapter dimers. (F) Barplot showing average % adapter dimers reads (n = 3) in LIDAR libraries from ESC total RNA. (G) Average % of reads preceded by their UMI sequence (with 0, 1, or 2 mismatches) in their genomic context. Data from LIDAR libraries starting from ESC total RNA (n = 3). (H) Barplot showing average (n = 3, ⊠ SEM) % of reads mapping to rRNA in LIDAR and ligation-based libraries from total ESC RNA. (I) Denaturing poly acrylamide-urea (9%) electrophoresis of total, < 200 nts, and < 50 nts RNA fractions isolated from ESC. (J) Average (n =3) biotype distribution of ESC non-coding RNAs, expressed as % of mapped reads, in LIDAR and ligation libraries from < 200 nts and < 50 nts ESC RNA inputs. (K) Barplot showing average number (n = 3, ⊠ SEM) of detected (with reads per million > 1) snoRNAs (left panel), or miRNA (right panel), with each sample subsampled to 1 million counts.

**Figure S2.**
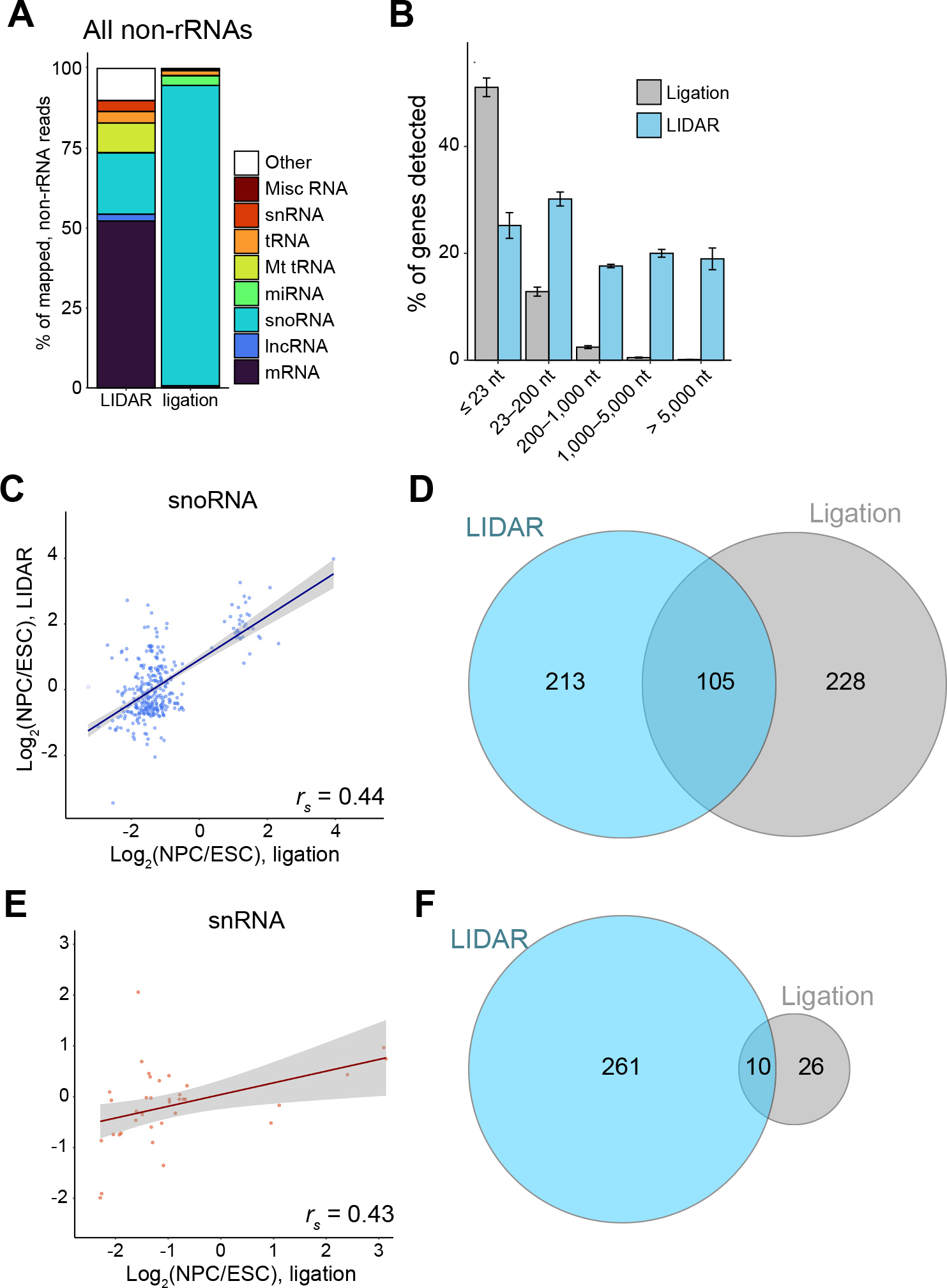
**Additional analyses on different RNA biotypes captured by LIDAR** (A) Average (n = 3) distribution of ESC non-rRNA biotypes, expressed as % of mapped reads, in LIDAR and ligation libraries from total ESC RNA. (B) Barplot of average percentage (n = 3, ⊠ SEM) of expressed ESC genes detected by LIDAR (blue) or ligation-based (grey) according to their size. Total ESC RNA used as input. All genes with mean TPM > 1 among all samples were included, and genes with TPM > 1 were considered expressed. (C) Correlation of log2-fold changes (NPC vs ESC) of differentially expressed snoRNAs detected by LIDAR and ligation. rs, Spearman’s rank correlation coefficient. (D) Venn diagram depicting overlap between differentially expressed snoRNAs detected in LIDAR (blue) or ligation-based libraries (grey). (E) Correlation of log2-fold changes (NPC vs ESC) of differentially expressed snRNAs detected by LIDAR and ligation. rs, Spearman’s rank correlation coefficient. (F) Venn diagram depicting overlap between differentially expressed snRNAs detected in LIDAR (blue) or ligation-based libraries (grey).

**Figure S3.**
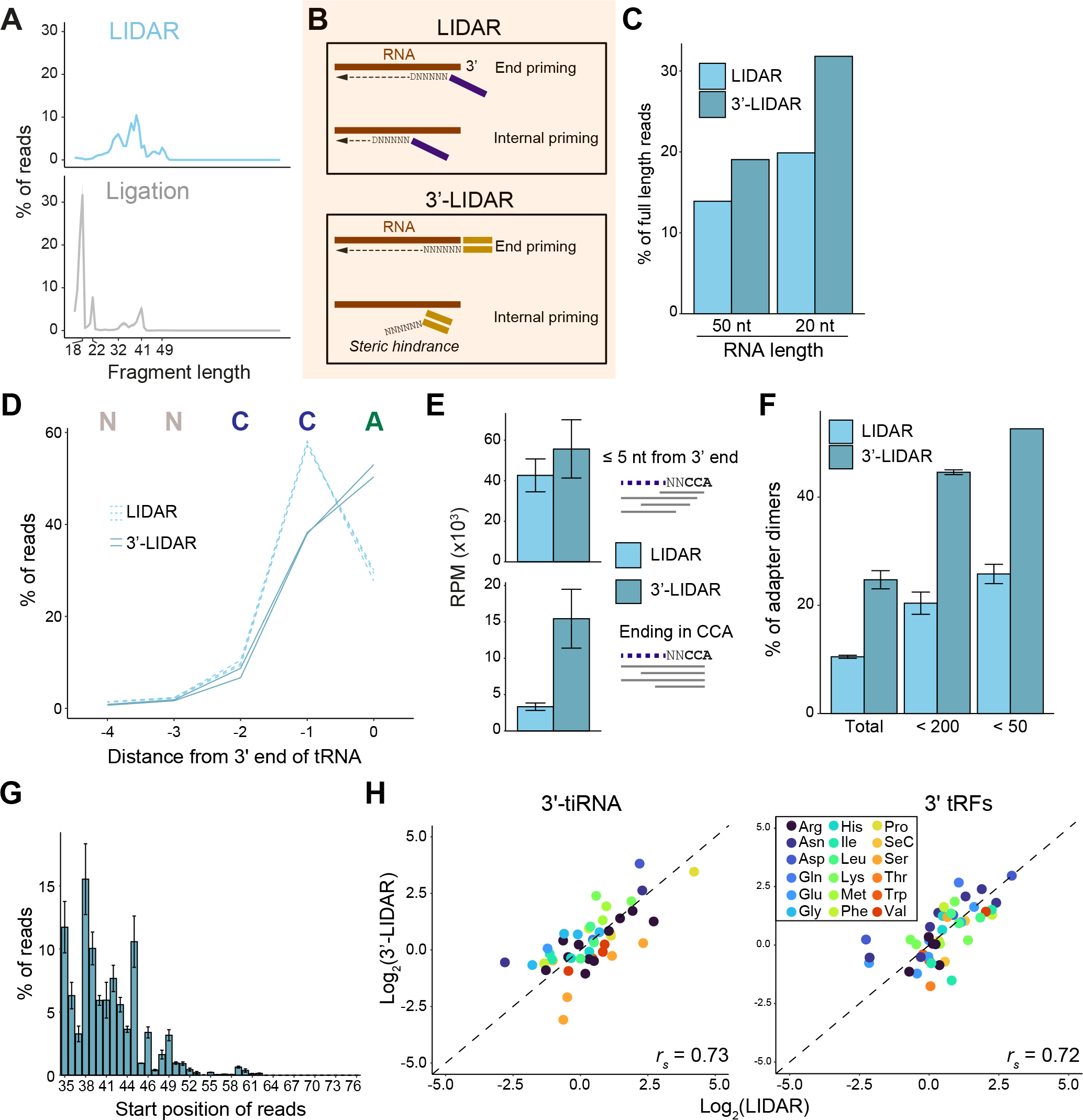
**Comparison of LIDAR and 3’-LIDAR** (A) Average size distribution of all reads mapping to 3’ tDRs (tiRNA or tRF), expressed as % of 3’ tDR reads, in LIDAR libraries from ESC RNA < 200nt (blue, top) or ligation-based libraries (grey, bottom). Data from 3 biological replicates. (B) Schematic representation of differences between LIDAR and 3’-LIDAR priming. The structure of the 3’-LIDAR RT primer is designed to inhibit internal priming, resulting in favored priming from the 3’. (C) Barplot showing percentage of end-to-end (full-length) reads mapping to synthetic 20 nts or 50 nts RNAs in LIDAR (light blue) or LIDAR-3 (dark blue) libraries. (D) Line plot showing % of tRNA reads mapping to the last 5 nucleotides at the 3’ end of mature tRNAs in LIDAR and 3’-LIDAR libraries from ESC RNA < 200 nts. Individual replicates are shown. (E) Average number of reads per million (RPM) (n = 3, ⊠ SEM) of reads ending within 5 nts (top panel) or at the CCA (bottom panel) of the 3’ end of mature tRNAs in LIDAR and 3’-LIDAR libraries from ESC RNA < 200 nts. (F) Average % of adapter dimers (n = 3 ⊠ SEM) in LIDAR (light blue) and 3’-LIDAR (dark blue) libraries. (G) Histogram of average (n = 3, ⊠ SEM) read beginning position frequency, expressed as % of all reads mapping to the corresponding tDR type, for 3’ tDRs (tRF or tiRNA) in 3’-LIDAR libraries starting from ESC RNA < 200 nts. (H) Scatterplot showing correlation of 3’-tRF (left) or 3’-tiRNA (right) iso-acceptors average log2 frequency (n = 3, ⊠ SEM), calculated over all reads mapping to 3’ tDRs, between LIDAR and 3’-LIDAR. rs, Spearman’s rank correlation coefficient.

**Figure S4.**
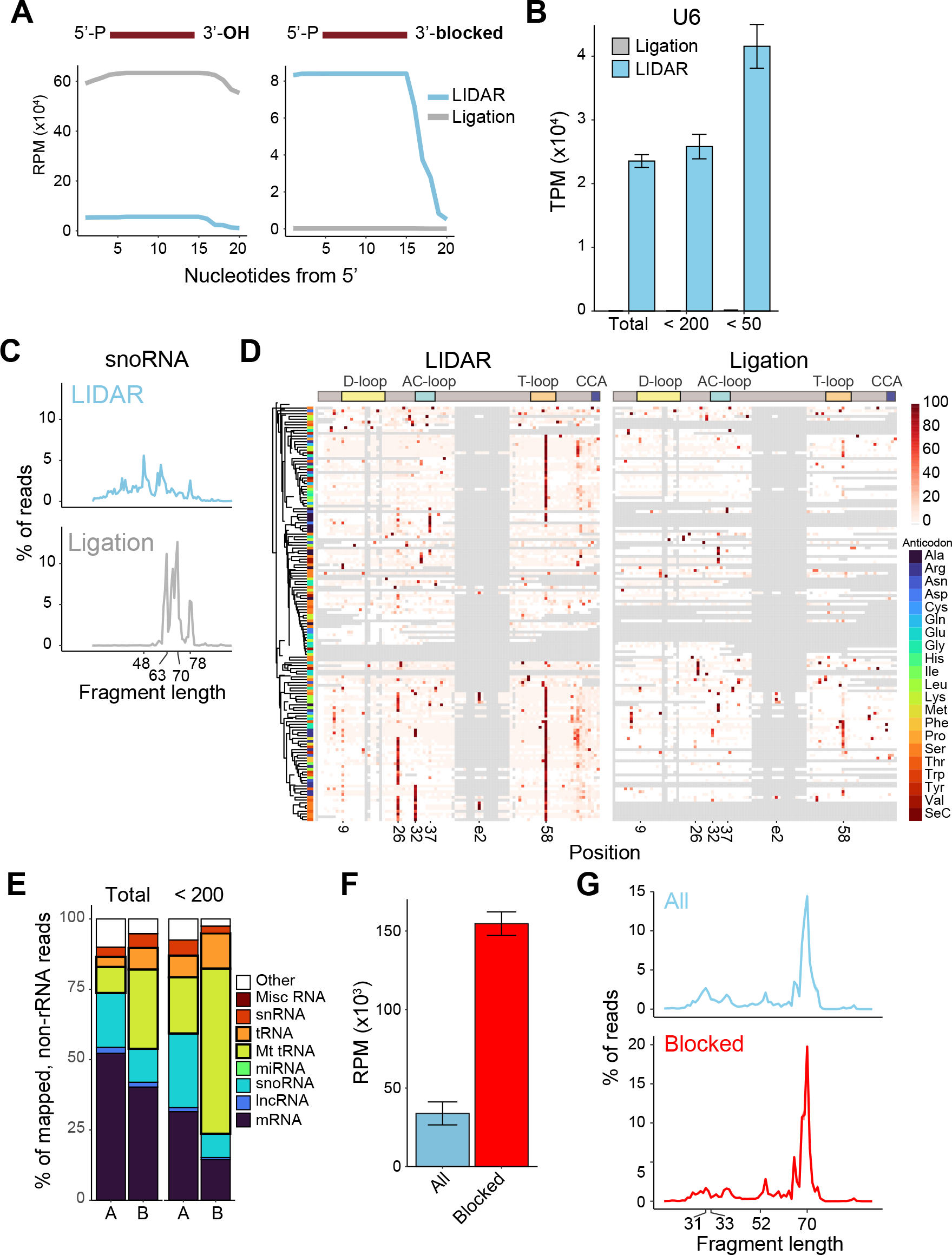
**Additional analyses on LIDAR libraries enriched for blocked RNAs** (A) Library coverage, expressed as reads per million (RPM) of synthetic 20 nts RNAs with 3’OH (left) or 3’ biotin (right) from LIDAR (blue) or ligation (grey) libraries. (B) Barplot showing average (n = 3, ⊠ SEM) transcripts per million (TPM) on U6 snRNA in LIDAR and ligation-based libraries from total, < 200nt, and < 50nt ESC RNA inputs. (C) Average size distribution of reads mapping to snoRNAs in LIDAR or ligation-based libraries starting from < 200 nts ¬¬ESC RNA. Data from 3 biological replicates. (D) Heatmap of average misincorporation rate (expressed as % detected) for every canonical position (column) tRNA iso-acceptor (rows) in LIDAR (left) or ligation-based (right) libraries starting from < 200 nts ESC RNA (n = 3). Gray indicates coverage = 0. All reads mapping to tRNAs were considered. (E) Average biotype distribution of non-rRNA genes, expressed as % of mapped reads, in LIDAR from all (A, n = 3) or blocked (B, n = 2) total and ESC RNA < 200 nts. (F) Barplot of reads mapping to full length tRNAs in LIDAR from all (n = 3) or blocked (n = 2) ESC RNA < 200 nts. (G) Average size distribution of all reads mapping to tRNAs, expressed as a % of all tRNA reads, in LIDAR libraries starting from all (n = 3) or blocked (n = 2) ESC RNA < 200 nts.

**Figure S5.**
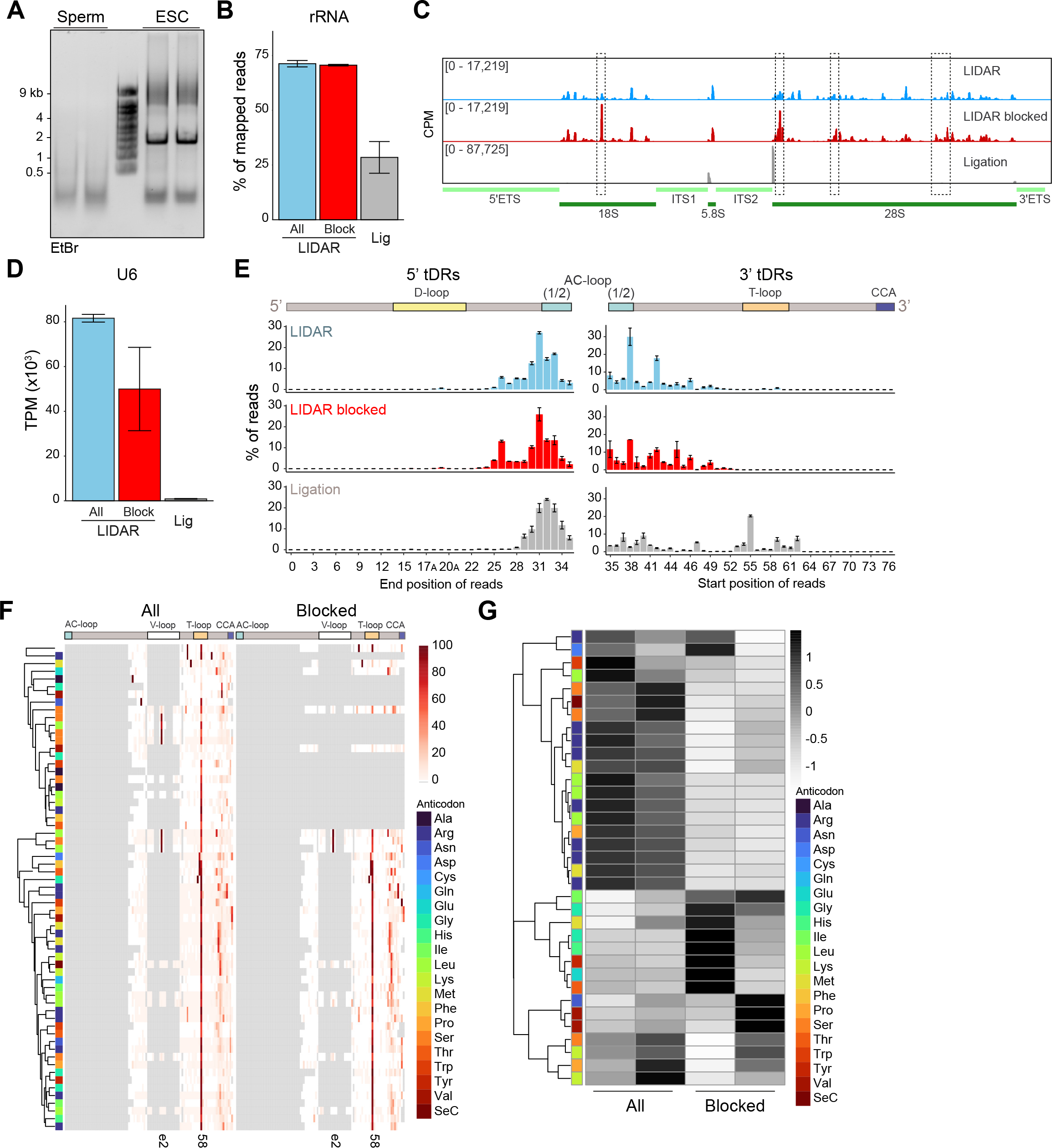
**Additional analyses on the RNA payload of mouse sperm** (A) Denaturing agarose gel electrophoresis analysis of total RNA from sperm and ESC (n = 2). (B) Reads mapping to rRNA in LIDAR and ligation-based libraries from all and blocked sperm RNA. Bars indicate the mean ± SEM (n = 2). (C) Read density on rRNA gene models in LIDAR libraries from all or blocked sperm RNA as well as ligation-based libraries. (D) Reads mapping to U6 in LIDAR from all or blocked sperm RNA and ligation-based-libraries. Bars indicate mean TPM ± SEM (n = 3). (E) Histogram of average (n = 3, ⊠ SEM) read end (left) or beginning (right) position frequency, expressed as % of all reads mapping to the corresponding tDR type, for 5’ tDRs (tRF or tiRNA) (left panel) and 3’ tDRs (tRF or tiRNA) (right panel) in LIDAR libraries starting from all (blue) or blocked (red) sperm RNA and ligation-based. Bars indicate the mean ⊠ SEM (n = 2). (F) Misincorporation rate (expressed as % detected) for every canonical position on 3’ tRNA fragment iso-acceptor in LIDAR libraries from all or blocked sperm RNA. Gray indicates coverage = 0. Data are from 2 biological replicates. (G) Heatmap of average (n = 2) log2 isoacceptor frequency, calculated over all reads mapping to 3’ tRNA fragments, of 3’ tRNA fragments in LIDAR libraries starting from all or blocked sperm RNA.

